# *Caenorhabditis elegans* HLH-30/TFEB has age-dependent effects on the transcriptome and is required for normal progeny number and larval survival under basal growth conditions

**DOI:** 10.64898/2026.11.19.767989

**Authors:** Ruiling Zhong, Sherlyn P. Wijaya, Ray A. O. Sinurat, Serenity D. Her, Qiyue Ying, Claire E. Richardson

## Abstract

**Background:** The transcription factor TFEB/HLH-30 is a well-established regulator of autophagy and lysosomal biogenesis during starvation and stress, and it plays important roles in development and homeostasis. Our previous work demonstrated that *Caenorhabditis elegans* HLH-30 also functions under basal physiological conditions in mature neurons, where it promotes the expansion of endo-lysosomal degradative capacity during early adulthood. These findings raised the question of whether HLH-30 promotes lysosomal gene expression more broadly during early adulthood, and, more generally, whether its basal transcriptional functions depend on animal age. To address these questions, we analyzed the impact of *hlh-30* loss on the transcriptome at two larval stages, L1 and L4, and two adult ages, Day 3 and Day 6, in hermaphrodites grown under basal conditions.

**Results:** We identified modest effects of *hlh-30* loss on the transcriptome that varied by stage/age. Enrichment analyses of genes downregulated in *hlh-30* mutants identified functions related to defense response at L1 and in adulthood, and to cuticle development in adulthood. On the other hand, a role for *hlh-30* in the basal expression of autophagy genes was exclusively detected in L1 larvae. A subset of genes encoding cathepsins showed decreased expression in the adult *hlh-30* mutant; however, our results indicate that HLH-30 is not broadly required for basal expression of most lysosome biogenesis genes in adulthood. We also detected a trend toward reduced expression of genes involved in germline function at the L4 stage, when spermatogenesis occurs, and found that loss of *hlh-30* resulted in reduced sperm and progeny numbers. In addition, we found that *hlh-30* is required for full progeny survival through larval development. Finally, we identified a gene with increased expression at all four stages in the *hlh-30* mutant, the germline transcription factor-encoding *ztf-28.* Knockdown of *ztf-28* partially suppressed the reductions in progeny number and survival, suggesting that its increased expression contributes to these phenotypes.

**Conclusions:** This study reveals roles for HLH-30 in transcriptome regulation under basal growth conditions that vary in both magnitude and content by animal age. Further, it demonstrates a previously unappreciated requirement for HLH-30 in promoting progeny number and survival, broadening our understanding of the physiological roles of this conserved transcription factor.

## Introduction

To maintain organismal health, cells require robust mechanisms of cellular homeostasis, including the coordination of protein quality control, metabolism, and stress responses [1, 2]. Transcriptional regulation allows cells to adjust these processes according to changing cellular demands [3, 4].

The MiT/TFE family transcription factors TFEB and TFE3 are well-established regulators of autophagy and lysosomal biogenesis, and they play important roles in maintaining cellular homeostasis [5–8]. Their activities are best understood in the context of stress and nutrient sensing: in nutrient-rich or non-stressed conditions, phosphorylation promotes cytoplasmic retention of TFEB and TFE3, whereas starvation and other cellular stresses promote their nuclear localization and induction of endosomal, lysosomal and autophagy-related genes [7, 9–12]. Beyond their canonical roles, TFEB and TFE3 are involved in several other physiological processes, including lipid metabolism, DNA damage response, embryonic development, and cell fate decisions, functioning broadly in coordinating cellular and organismal homeostasis [13–20].

*C. elegans* HLH-30, the sole homolog of TFEB and TFE3, likewise regulates lysosomal and autophagy-related gene expression and promotes organismal stress responses [21–23]. Starvation induces HLH-30 nuclear localization and transcriptional activity, promoting autophagy, lipid metabolism, and survival during nutrient deprivation [22, 24–27]. HLH-30 also promotes survival and recovery during starvation-induced developmental arrest, including L1 arrest and adult reproductive diapause [28–30]. These functions extend beyond autophagy and lysosomal biogenesis: HLH-30 regulates chromatin reorganization and cell quiescence during L1 starvation [29], and it promotes germline stem cell resilience during adult reproductive diapause through regulation of growth signaling, largely independently of canonical autophagy [28]. In addition, HLH-30 regulates innate immunity, resistance to other stresses, and longevity [21, 23, 31–34]. Together, these studies establish HLH-30 as an important regulator of organismal physiology, although its functions have been characterized predominantly in the context of environmental or physiological stress.

Our previous work demonstrated that HLH-30 also functions under basal physiological conditions in mature neurons, where it promotes the expansion of endo-lysosomal degradative capacity during early adulthood and is required for adult neuronal maintenance [35]. Notably, basal HLH-30 activity was sufficient to promote this expansion without nuclear enrichment associated with stress-induced activation. We also found that neuronal lysosomal function declines during mid-adulthood, a phenomenon that likewise occurs in mammalian neurons, and that decline is accompanied by reduced HLH-30-dependent transcription [35, 36]. These findings raised the question of whether HLH-30 promotes a similar expansion of lysosomal capacity more broadly during early adulthood, and whether its basal transcriptional functions vary with animal stage/age. More generally, the extent to which HLH-30 regulates gene expression and organismal physiology across different ages under normal growth conditions remains poorly understood.

Here, we investigated the role of HLH-30 in regulating gene expression and organismal homeostasis across different life stages under basal conditions. Transcriptomic profiling from the fourth larval stage (L4) to mid-adulthood revealed age-dependent effects of HLH-30 on genes involved in defense responses, cuticle production, and reproduction. Functional analyses further revealed that HLH-30 is required for reproductive fitness and progeny survival under non-starved conditions. We identified increased expression of transcription factor *ztf-28* as a contributor to the reproductive defects of *hlh-30* mutants. Additional transcriptomic profiling at L1 identified a prominent role for HLH-30 in promoting basal expression of autophagy- and lysosome-related genes during early larval development. Together, these findings reveal previously unrecognized roles for HLH-30 in reproductive fitness and progeny survival and demonstrate that its transcriptional functions under basal conditions vary across developmental stages.

## Methods

### *C. elegans* strains and maintenance

For maintenance, *C. elegans* were grown on nematode growth medium (NGM) seeded with *E. coli* OP50 at room temperature (22 ℃). Adult worm age is defined by the number of days starting from the L4 stage (Day 0). Unless otherwise indicated, all experiments were performed at 20 ℃. Wild-type N2, JIN1375 *hlh-30(tm1978) IV*, OH17942 *tts-1(ot1234) X*, and RW12281 *ztf-28(st12281) V* strains were obtained from the *Caenorhabditis* Genetics Center (CGC), which is funded by NIH Office of Research Infrastructure Programs (P40 OD010440). The *cpr-4(syb11759[7xGFP11::NLS::PEST])* allele was generated at SunyBiotech (Fuzhou, Fujian, China). Strains used in this study are listed in Additional file 8 - Supplementary Table 1.

### RNA library preparation and sequencing

Total RNA was extracted from L1, L4, Day 3, and Day 6 WT and *hlh-30(tm1978)* hermaphrodites at least three times independently as described [37]. For generating synchronized L1 larvae, embryos were isolated from gravid adults using an alkaline hypochlorite (“bleach”) solution, and then they were hatched in M9 solution for 2-5 hours. For adult samples, L4 worms were transferred to 51 µM fluorodeoxyuridine (FuDR) plates, and they were transferred to new plates every other day. Total RNA was isolated using the Trizol method [37]. cDNA libraries were prepared for Illumina RNA sequencing and 3’ end counting using the Plasmidsaurus platform (Plasmidsaurus, Louisville, KY, USA).

### RNA sequencing and read processing

Libraries were sequenced by Plasmidsaurus (3′-end RNA-seq, single-end reads with unique molecular identifiers). Plasmidsaurus filtered the reads with fastp [38], aligned them to the *C. elegans* genome (WBcel235) with STAR [39] and removed PCR duplicates by UMI-based deduplication. Gene counts were generated from the deduplicated alignments with featureCounts [40] using exon features of the Ensembl release 115 annotation [41]. Reads were counted forward-stranded, as 3′-end library reads match the sense strand of the transcript. Genes with fewer than 10 reads summed over all samples were removed. Software versions and quality control steps are listed in Additional file 9 - Supplementary Methods.

### Differential expression

The full set of RNA-seq data generated for this study had 34 samples of wild type and *hlh-30(tm1978)*, with three biological replicates per genotype at L1 and L4, six at Day 3 of adulthood, and five at Day 6 of adulthood. For three of the adult biological replicates, the animals were grown at 20 °C, and for the rest of the adult samples, the animals were grown at room temperature (22 °C). A model that included the temperature variable found no expressed gene that differed between the temperatures, and the temperatures did not separate in the adult PCA (Additional file 7 - Fig. S1C). Based on these results, the samples from 20 °C and room temperature were combined for all subsequent analysis. Two subsets were also analyzed, L4 to Day 6 (28 samples) and Day 3 and Day 6 (22 samples, for the adult-only principal component analysis and clustering). For each sample set, one factor of unwanted variation (W_1) was estimated with the RUVs method of RUVSeq [42], using the replicates of each genotype and stage as control sample groups. Across all 34 samples, 21,045 genes were detected. Differential expression was tested with DESeq2 [43] in R 4.5.2 [44] with the design ∼ W_1 + replicate + condition, where condition combines genotype and stage. Wald tests were run between stages within each genotype (developmental transitions) and between genotypes at each stage. Log2 fold changes were shrunk with ashr [45], and P values were adjusted with the Benjamini-Hochberg method [46]. Genes with an adjusted P < 0.05 and an absolute log2 fold change ≥ 1 were called differentially expressed genes (DEGs). Some analyses here and in the following sections were performed using WormIE (https://wormie.bio) [47].

### Expression visualization and gene clustering

For principal component analyses, sample correlation heat maps and clustering, counts were variance-stabilized with DESeq2, and the replicate effect and W_1 were removed with limma removeBatchEffect [48] while keeping genotype and stage differences. Principal component analyses used the 500 most variable genes. Sample correlation heat maps used Pearson correlation of the 1,000 most variable genes across the genotype and stage means, ordered by average-linkage hierarchical clustering.

For gene clustering, the mean expression of each gene in each genotype and stage was z-scored across the genotype and stage means, and the DEGs of all contrasts in a sample set were clustered on these z-scores. k-means clustering was run with 25 random starts, and the number of clusters was chosen by the elbow method (k = 6 for the full set and the L4 to Day 6 subset, k = 4 for the Day 3 and Day 6 subset). As a density-based comparison, genes were also clustered with HDBSCAN in the R package dbscan [49]. HDBSCAN assigned 523 genes, 10% of the total DEGs, as noise and produced relatively few interpretable clusters, making the k-means clustering more useful for further analysis. Clusters were displayed with UMAP in the R package uwot [50].

### Gene Ontology analysis

Gene Ontology (GO) enrichment [51] was tested with clusterProfiler [52] for biological process, cellular component and molecular function, with all genes that passed the count filter as background. Terms with a Benjamini-Hochberg adjusted P < 0.05 were considered enriched. Gene lists tested were the k-means clusters, the up- and down-regulated DEGs between genotypes at each stage and the gene lists from the comparison of developmental changes between genotypes (next section). The terms shown in the figures were selected from the enriched terms of each list.

### Comparison of developmental expression changes between genotypes

For each developmental transition assessed (L1 to L4, L4 to Day 3, L4 to Day 6 and Day 3 to Day 6), the DEGs of wild type and *hlh-30(tm1978)* were compared. A low-count gene that DESeq2’s independent filtering removed from at least one of the two genotypes’ contrasts was left out, so that it was not counted as exclusive to the other genotype. A gene was "exclusive" when it was a DEG in only one genotype, "shared" when it was a DEG in both genotypes with an absolute difference in log2 fold change < 1, and "differential" when that difference was ≥ 1. Differential genes with fold changes of opposite sign were called "opposite". Differential genes that changed in the same direction in both genotypes were called "more up" or "more down" in the genotype with the larger change.

For Additional file 7 - Fig. S3, two gene sets were taken from the GO terms enriched in these gene lists for the three adult transitions. The oocyte and germline set was the 31 genes of the terms that name a germline process, and the sperm set was the 18 genes of the pseudopodium term, which holds the major sperm protein genes. Expression was z-scored per gene across the genotype and stage means. Differences between genotypes at each stage were tested for each set with limma camera [53] and for each gene with the DESeq2 contrast of that stage.

### Fluorescence confocal microscopy

Confocal imaging was performed as described [35]. Worms were transferred to a slide with a 3% agarose pad made by melting agarose in M9 buffer. Worms were immobilized in 10 mM levamisole in M9 buffer. Images were captured using a Nikon eclipse Ti2 microscope paired with a CSU W1 SoRa confocal scanner unit and a YOKOGAWA spinning disc unit with a Plan Apo VC 60xA/1.20 WI objective. Image settings (laser power and exposure time) were identical for all genotypes and treatment groups across experiments. Images were taken with 0.5 μm step size in z-stack.

### Fecundity assay

Fecundity assays were performed at 20 °C using 5-6 animals per genotype/treatment/condition as replicates. For each replicate, one L4 worm (P0) was picked onto one plate and transferred onto a fresh plate every day until reproduction ceased (Fig. 5A). The number of progeny laid, including unhatched eggs and hatched L1s, was counted after the P0 was transferred each day. The number of unfertilized oocytes was also counted at the same time. The plates were left at 20 ℃ for two more days and the number of L4 larvae and adult progeny was counted. Because of manual counting errors, the number of total L4 + adult progeny counted was sometimes greater than the number of eggs + L1s counted on a plate; when that occurred, the number of eggs + L1s was corrected to match the number of L4s + adults. The number of total progeny laid and the number of total adult progeny were calculated as the sum for one worm over its lifespan. The survival rate was calculated as the number of total adult progeny divided by the number of total progeny laid. All fecundity assays were blinded except for the *ztf-28* RNAi experiments. All fecundity assays were performed in at least two independent experimental trials that were combined for analysis.

### Auxin treatment

L4 worms were transferred to 1 mM naphthaleneacetic acid (K-NAA) (PhytoTech Labs, Lenexa, KS, USA) plates seeded with *E. coli* OP50. For the fecundity assay to determine tissue specificity of HLH-30, L4 worms were picked in the subsequent generation, and auxin treatment was continued throughout the experiment.

### DAPI staining and analysis

DAPI staining was performed as described [54]. Briefly, worms were synchronized by the bleach method, hatched overnight in M9 media, and dropped onto NGM plates seeded with *E. coli* OP50. Worms were harvested at Day 1 and fixed in 95% ethanol for 10 min. Ethanol was then removed, and 50 μL of Vectashield mounting medium with DAPI (Vector Laboratories, Burlingame, CA, USA) were added to the samples. Samples were stored in the dark at room temperature, and they were mounted on slides without spacers for imaging.

Quantification of germ cells was performed as described, counting the number of germ cell nuclei along a lengthwise line through the gonad [55]. Sperm were counted in the gonad arm closer to the cover slip by drawing an ROI around the sperm, which were predominantly but not exclusively in the spermatheca based on position [56]. Sperm were identified by their distinctively small nuclei. To facilitate counting, non-sperm DAPI-stained nuclei in the region were deleted manually from z-slices. Then, a z-projection that included all the sperm was generated. Sperm were roughly selected using “find maxima;” then, non-sperm maxima were manually removed, and sperm that were not automatically identified were manually added to get the complete count. Animals in which too many non-sperm nuclei were intermixed with the sperm to reasonably remove were not counted. Of 39 animals counted, 24 were scored blind to genotype (10 of 19 wild-type and 14 of 20 *hlh-30* mutant). There was no substantial difference in the sperm counts scored blind versus not: for WT, scored-blind mean = 174.5 ± 14.5 SEM, not-blind mean = 176.6 ± 10.6 SEM; for the *hlh-30* mutant, scored blind mean = 136.3 ± 7.3 SEM, not-blind mean = 132.5 ± 9.1 SEM.

### RNAi treatment

L4 worms were transferred to 1 mM isopropyl β-D-1-thiogalactopyranoside (IPTG) plates seeded with *E. coli* HT115 containing empty vectors L4440, *unc-22* RNAi or *ztf-28* RNAi clones from the Ahringer Library [57]. L4 worms were picked from the subsequent generation for the fecundity assay, and the RNAi feeding was continued throughout the experiment.

### Analysis of microscopy data

Images were analyzed using ImageJ software. To analyze the real-time *cpr-4* transcriptional reporter, a maximum projection image was made from each z-stack, an ROI was drawn around each worm, and the maximum fluorescence intensity value, which arose from an intestinal nucleus, was recorded. To analyze *tts-1p::gfp* expression, a maximum projection image was made from each z-stack. An ROI was drawn around each worm, the mean GFP intensity was recorded, and then the ROI was moved to an area of the image with no worms to obtain the background mean value. The mean intensity per worm was calculated by subtracting each background mean from its corresponding worm mean. To analyze ZTF-28::GFP expression, the germ cells in the gonad arm closer to the cover slip for each worm were used. Three 12 μm × 12 μm ROIs were drawn over different regions of the germline per worm. For each ROI, the z-slice with the highest mean fluorescence intensity value within the germline was used, and the mean intensity was recorded. The three values per animal were averaged to generate one data point per animal.

### Statistical analysis

Statistical analyses were performed in either Microsoft Excel or GraphPad Prism 10. Single parametric pairwise comparisons were analyzed using unpaired two-tailed Student’s *t* test, and multiple parametric comparisons were analyzed using one-way analysis of variance (ANOVA) test with Tukey post-test or two-way ANOVA with Sidak post-test as indicated in the figure captions. The *cpr-4, tts-1,* and *ztf-28* fluorescence intensity data were log-transformed for statistical analysis due to heteroscedasticity of the data.

### Motif analysis

Promoters (2,000 bp upstream to 200 bp downstream of the transcription start site) were scanned with motifmatchr [58] for the HLH-30 motif from JASPAR (MA1449.1) [59] at a match P < 1e-4, so that sites include close variants of its consensus TCACGTGA. For each DEG list, the fraction of genes with at least one site was compared with the genes that passed DESeq2’s independent filtering in at least one mutant vs wild-type contrast by a two-sided Fisher’s exact test with Benjamini-Hochberg correction over the 12 tests. In the promoter schematics, E-box sites are exact CACGTG matches, the core of the CLEAR element [5]. In the *tts-1*, *cpr-4* and *ztf-28* promoters, we also counted the DAF-16-binding element (DBE; TRTTTAC and TTGTTTAC) and the PQM-1-bound DAF-16-associated element (DAE; TGTAAG and TGATAAG), which are enriched at HLH-30 binding sites [34].

Enrichment of all transcription factor motifs was tested with SEA [60, 61] in the MEME Suite v5.5.7 [61], as in Chang et al. [62], with the motifs of JASPAR 2022 nematodes [63] and CIS-BP 2.00 *C. elegans* [64], in the 500 bp and the 2,000 bp upstream of the TSS. The control sequences for the DEGs of each stage were the promoters of the other tested genes at that stage (adjusted P ≥ 0.05), and for the DEGs of any stage, the promoters of genes with adjusted P ≥ 0.05 at every stage. Motifs with q < 0.05 were considered enriched when the gene of their transcription factor was expressed at that stage.

### Analysis of HLH-30 ChIP-seq data

HLH-30::GFP ChIP-seq data from the modENCODE project [65] for L4 larvae (GEO GSE48751) and late embryos (GEO GSE48752) were obtained from ChIP-Atlas [66] as MACS2 peaks (Q < 1e-5) and coverage tracks aligned to ce11. For the metagene profile, ChIP coverage was divided by input coverage in bins over ±3 kb around the transcription start sites of protein-coding genes, and each profile was scaled so that the flanks beyond ±2.5 kb equal 1. L4 peaks were taken as reproducible when the summits of the two replicates were within 100 bp.

### HLH-30 binding and published gene lists

The analyses of HLH-30 binding, motifs and published gene lists used the DEGs of the full sample set. A gene was counted as bound by HLH-30 when a reproducible L4 ChIP-seq summit was within 1,000 bp upstream to 500 bp downstream of its TSS. The bound share of each DEG list was compared with that of the other genes by a two-sided Fisher’s exact test, and between the DEG lists of two stages by an exact permutation test that allows a gene to be in both lists. Our DEGs were compared with published HLH-30 gene lists (sources in Additional file 9 - Supplementary Methods) by a two-sided Fisher’s exact test. Core autophagy and lysosome genes were taken from Zhang and Meléndez [67] and Sun et al. [68]. P values were adjusted with the Benjamini-Hochberg method.

### Use of large language models

Claude Fable 5.1 (claude-fable-5-1) and Claude Opus 5.5 (claude-opus-5-5), both from Anthropic, were used to refine the writing and the code. ChatGPT (OpenAI) was used to assist with literature searches, blinding microscopy images, and manuscript editing.

## Results

### Loss of *hlh-30* causes age-dependent changes in gene expression under basal conditions

To investigate *hlh-30*-dependent gene expression for adult maintenance under basal conditions, we performed bulk RNA-seq on wild-type (WT: N2) and the *hlh-30(tm1978)* mutant at the L4 larval stage, and at Day 3 and Day 6 of adulthood (Fig. 1A). The *hlh-30(tm1978)* allele is a frameshifting deletion and putative null allele. Principal component analysis (PCA) and Pearson correlation analysis followed by hierarchical clustering showed that the samples clustered primarily by age and then by genotype, with reproducibility among replicates (Fig. 1B-D). Day 3 and Day 6 samples clustered more closely with each other than with L4 samples, indicating substantial transcriptomic changes between the final larval stage and early adulthood in both WT and *hlh-30* mutant worms (Fig. 1B-C). PCA with only the adult samples showed that PC1 was associated with age and PC2 was associated with loss of *hlh-30* (Fig. 1D).

**Fig. 1.**
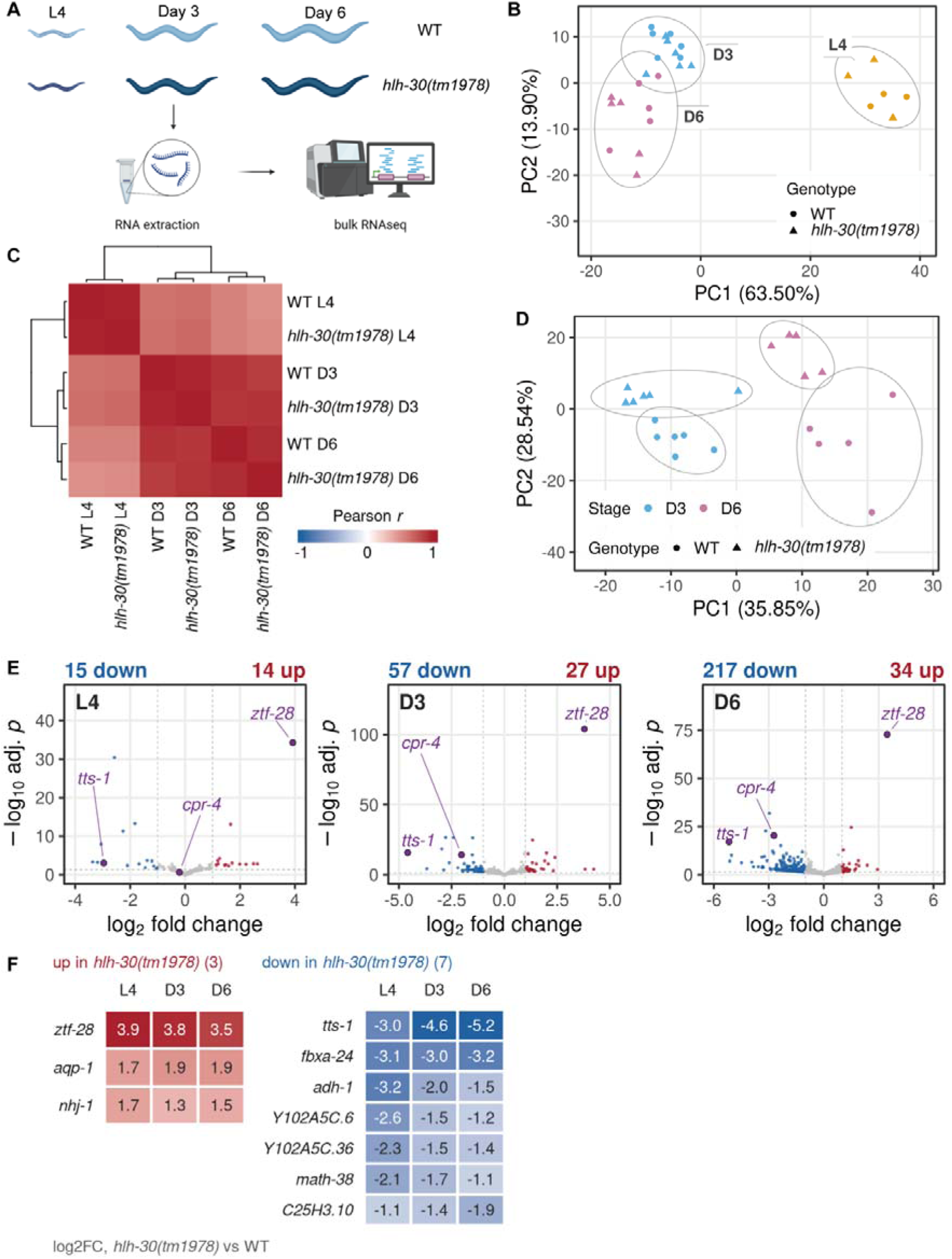
The *hlh-30* mutant shows age-dependent changes in gene expression under basal conditions. (A) Experimental workflow for bulk RNA-seq. (B-D) PCA (B, D) and hierarchical clustering (C) show grouping of samples by stage and genotypes. (E) Volcano plots showing differentially expressed genes (DEGs) between WT and the *hlh-30* mutants at each age. (F) DEGs shared across the three ages.

We performed pairwise comparisons between WT and the *hlh-30* mutant at each age using the differential gene expression analysis program DESeq2 [43]. Compared with WT at the same stages, 29, 84, and 251 genes were differentially expressed in the *hlh-30* mutant at L4, Day 3, and Day 6, respectively (Fig. 1E, Additional file 1). These *hlh-30*-dependent differentially expressed genes (DEGs) comprise 0.2%, 0.6%, and 1.8% of the expressed genes at these stages. Ten DEGs were shared among all stages, and another 47 were shared only between Day 3 and Day 6 (Fig. 1F, Additional file 7 - S1A). Thus, loss of *hlh-30* causes relatively modest transcriptional changes under basal conditions, with differences becoming more pronounced during adulthood.

To validate the transcriptomic analysis, we assessed the expression of fluorescent reporters for two genes that showed reduced expression in the *hlh-30* mutant: *cpr-4* and *tts-1* (Fig. 2). As *cpr-4* encodes cathepsin B, a lysosomal resident protein, it is representative of the lysosomal biogenesis genes that are canonically regulated by TFEB/HLH-30. We generated a real-time fluorescent transcriptional reporter for *cpr-4(syb11759),* wherein a destabilized nuclear GFP11::NLS::PEST is expressed from the endogenous *cpr-4* locus and reconstituted with ubiquitously expressed GFP1-10 [35]. The destabilizing PEST sequence targets the protein for rapid degradation, so the GFP fluorescence intensity reports the recent *cpr-4* transcription. We observed *cpr-4* reporter expression exclusively in the intestine at L4, Day 3, and Day 6 (Fig. 2A). The mean *cpr-4* reporter fluorescence intensity was greater in WT than in the *hlh-30* mutant at all three ages, though the difference was not statistically significant at Day 3 (Fig. 2A, C).

**Fig. 2.**
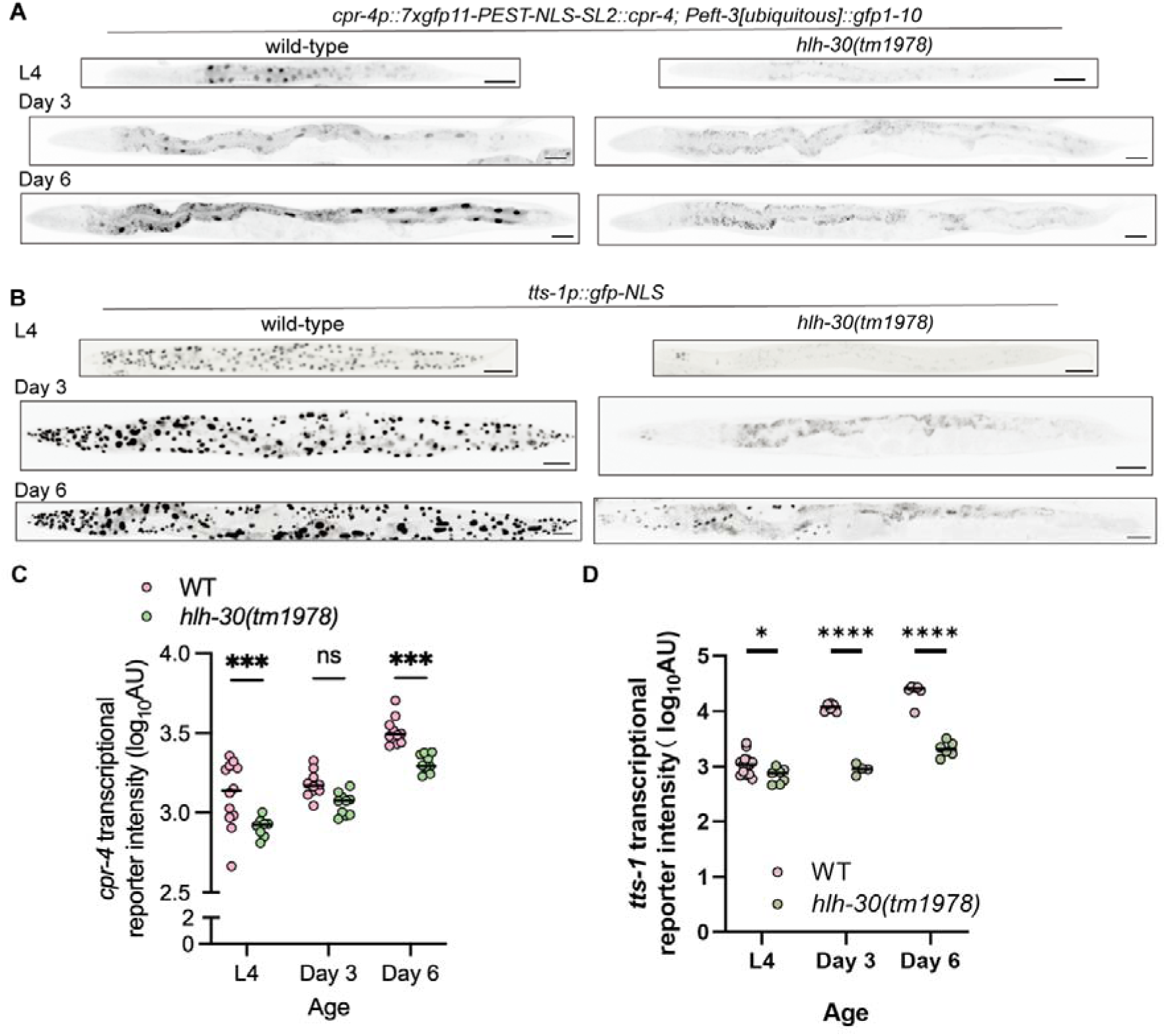
Fluorescent reporter validation of *hlh-30* mutant DEGs. (A-B) Example images of the real-time fluorescent reporter for *cpr-4* (A) and the fluorescent reporter for *tts-1* (B) in WT versus the *hlh-30* mutant. Scale = 50 μm. (C-D) Quantification of fluorescence intensity of the *cpr-4* (C) and *tts-1* (D) transcriptional reporters. Each data point is one worm. The y-axis is log transformed for visualization. ****P<0.0001, ***P<0.001, *P<0.05, ns: not significant; two-way ANOVA with Tukey post-test.

We next investigated the transcription level of the non-coding RNA *tts-1*, which was down-regulated in the *hlh-30* mutant at all stages (Fig. 1E). We used the *tts-1(ot1234)* allele, which co-transcriptionally drives GFP-tagged-histone from the endogenous *tts-1* promoter and shows broader expression compared with *cpr-4* (Fig. 2B). In the *hlh-30* mutant, the *tts-1* transcriptional reporter showed lower expression compared to WT at all three timepoints (Fig. 2B, D). These results broadly corroborate the RNA-seq results (Fig. 2B, D).

### Basal HLH-30 activity supports age-dependent expression of defense response and cuticle-related genes

To look more closely at the effects of HLH-30 on gene expression from the final larval stage to mid-adulthood, we performed hard (k-means) and soft (HDBSCAN) clustering on the 5,044 genes that were differentially expressed in pairwise comparisons of samples matched by either age or genotype [69–71]. Both methods detected the same two major transcriptional groups (Fig. 3A, Additional file 7 - S1B, Additional file 2). Because k-means identified six clusters with distinct temporal expression profiles, whereas HDBSCAN produced fewer interpretable clusters, we used k-means for further analysis (Fig. 3B). The expression levels of 176 genes gradually increased with age in WT but decreased in the *hlh-30* mutant (cluster 1), suggesting that HLH-30 is required for the age-associated induction of this transcriptional program. Gene Ontology (GO) enrichment analysis revealed enrichment for three Cellular Component terms: nucleosome, protein-DNA complex, and collagen trimer (Fig. 3C). Cluster 1 was also enriched for biological processes related to defense responses and cell killing, consistent with previous studies showing that HLH-30 promotes *C. elegans* innate immune responses and pathogen resistance (Fig. 3D) [23, 31, 33, 72–74]. Clusters 2-6, comprising 4,868 genes (97% of the total), exhibited pronounced changes between L4 and adulthood, whereas transcriptional differences between WT and the *hlh-30* mutant at each timepoint were relatively modest (Fig. 3B). Therefore, developmental and age-related changes were the dominant determinant of gene expression in this dataset.

**Fig. 3.**
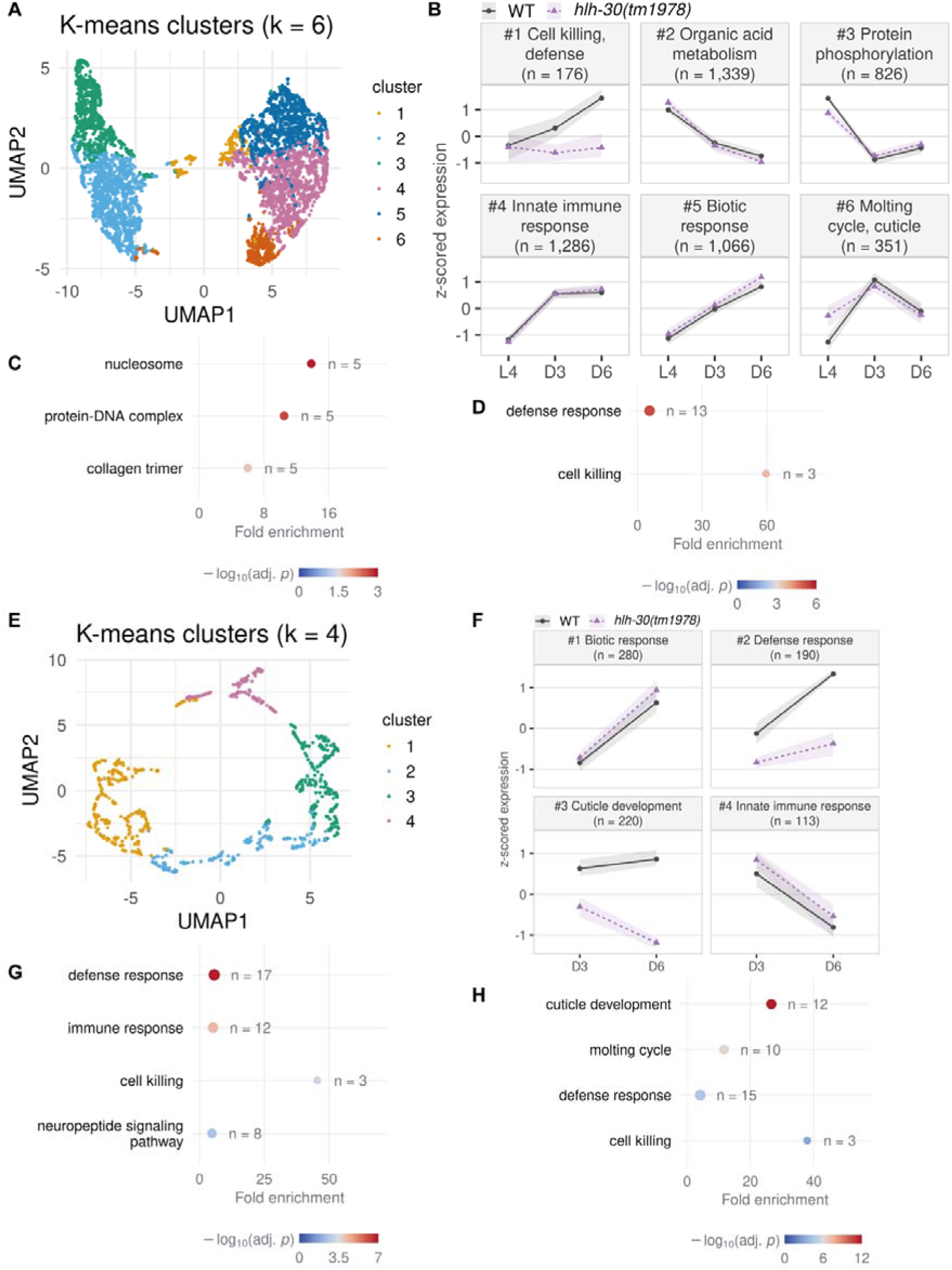
Clustering reveals age-dependent expression patterns in genes related to immune response. (A-B) K-means clustering of 5,044 DEGs among L4, Day 3, and Day 6 WT and *hlh-30*mutant samples identified six distinct temporal expression profiles. (C–D) Gene Ontology (GO) Cellular Component (C) and Biological Process (D) enrichment analyses of genes in Cluster 1 (176 genes), whose expression level increased with age in WT but decreased in the *hlh-30* mutant. (E–F) K-means clustering of 803 DEGs from the WT vs *hlh-30* mutant Day 3 and Day 6 samples identified four distinct adult expression profiles. (G–H) GO Biological Process enrichment analyses of genes in Clusters 2 (G) and 3 (H), which exhibited genotype-dependent expression trajectories during adulthood.

To further explore expression patterns during adulthood, we performed k-means clustering on 803 DEGs among the Day 3 and Day 6 samples, identifying four expression trajectories (Fig. 3E-F, Additional file 2). Clusters 1 and 4 primarily reflected age-related transcriptional changes shared between WT and the *hlh-30* mutant (Fig. 3F). We therefore focused on clusters 2 and 3, in which expression patterns differed between WT and the *hlh-30* mutant. Cluster 2 contained 190 genes whose expression levels were highly induced in WT from Day 3 to Day 6 but failed to undergo this age-associated increase in the *hlh-30* mutant. Cluster 3 comprised 220 genes that maintained fairly stable expression in WT across the two timepoints but showed lower expression in the *hlh-30* mutant at Day 3 with a further decline by Day 6. GO enrichment analysis of both clusters identified biological processes related to pathogen defense and cell killing (Fig. 3G-H). Consistent with the analysis including L4 samples, these enrichments support a role for basal HLH-30 activity in expression of innate immunity-related genes during adulthood, even in the absence of infection. Cluster 2 was also enriched for genes involved in neuropeptide signaling (Fig. 3G), and Cluster 3 for genes involved in cuticle formation (Fig. 3H).

To identify additional biological processes potentially affected by loss of *hlh-30*, we compared age-dependent gene expression changes between WT and the *hlh-30* mutant. We identified genes that were differentially expressed during the L4-to-Day 3, Day 3-to-Day 6, and L4-to-Day 6 transitions in one genotype but not the other, and performed GO enrichment analyses on these gene sets (Fig. 4, Additional file 7 – Fig. S2, Additional file 3). Since this analysis compares genes that pass differential-expression thresholds in each genotype, it provides a qualitative view of shared and genotype-specific transcriptional changes rather than a quantitative comparison.

**Fig. 4.**
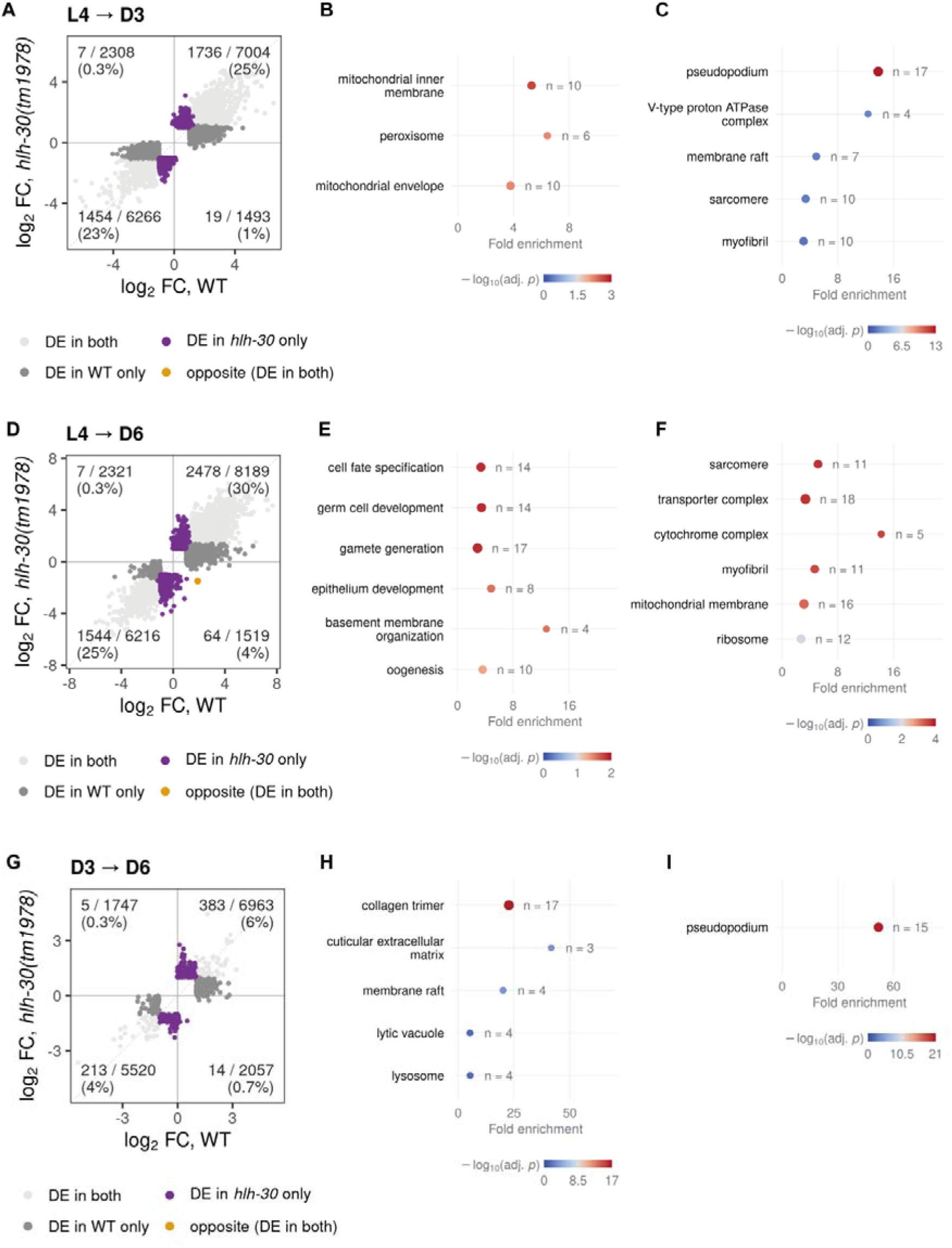
Comparative differential expression analysis suggests HLH-30-dependent transcriptional changes across stages. (A) Comparison of DEGs between WT and the *hlh-30* mutant during the L4-to-Day 3 transition. (B-C) GO cellular component enrichment analysis of genes that are exclusively down-regulated in the *hlh-30* mutant (B) and WT (C) during the L4-to-Day 3 transition. (D) Comparison of DEGs between WT and the *hlh-30* mutant during the L4-to-Day 6 transition. (E) GO biological process enrichment analysis of genes that are exclusively upregulated in the *hlh-30* mutant during the L4-to-Day 6 transition. (F) GO cellular component enrichment analysis of genes that are exclusively down-regulated in the *hlh-30* mutant during the L4-to-Day 6 transition. (G) Comparison of DEGs between WT and the *hlh-30* mutant during the Day 3-to-Day 6 transition. (H-I) GO cellular component enrichment analysis of genes that are exclusively downregulated in the *hlh-30* mutant (H) and upregulated in WT (I) during the Day 3-to-Day 6 transition.

Terms related to lysosome, mitochondria, and peroxisome were enriched for possible dependence on HLH-30 (Fig. 4B, H). We also identified GO terms related to the germline. For instance, the pseudopodium term, which in *C. elegans* mainly reflects major sperm protein genes, was enriched in genes that were exclusively downregulated from L4 to Day 3 in WT, and also in genes that were exclusively upregulated from Day 3 to Day 6 in WT (Fig. 4C, I). The downregulation of sperm genes between L4 and Day 3 is consistent with the hermaphrodite germline developmental timing, wherein the germline irreversibly switches from spermatogenesis to oogenesis around the L4-to-adult molt (Fig. 4C)[75]. In addition, germ cell development and gamete generation GO terms were enriched in genes that were exclusively upregulated in the *hlh-30* mutant between L4 and Day 6 (Fig. 4E). Taken together, these analyses suggest that loss of *hlh-30* may affect germ cell development.

As germ cell-related GO terms emerged from the comparative differential expression analysis, we examined the expression trajectories of these genes across the developmental time course, separately for oocyte and germline genes and for sperm genes (Additional file 7 - Fig. S3). On average, both gene sets showed a trend toward lower expression in the *hlh-30* mutant at the L4 stage compared to WT, and the oocyte and germline genes showed a trend toward higher expression in the *hlh-30* mutant in the adult timepoints compared to WT, though these differences were not statistically significant (Additional file 7 - Fig. S3A-D).

### HLH-30 is required for reproductive fitness under basal conditions

The comparative differential expression analysis suggested that HLH-30 may regulate reproductive physiology in L4 and adulthood. In addition, when maintaining *hlh-30* mutant worms, we anecdotally but consistently observed slower population expansion compared to WT strains. To test whether HLH-30 is required for reproduction, we performed a fecundity assay in which individual hermaphrodites (P0s) were transferred to fresh plates daily starting at L4 until reproduction stopped. For each P0, the number of progeny produced, the number of progeny that survived to the L4 stage, and the number of unfertilized oocytes were quantified throughout the reproductive period (Fig. 5A).

The *hlh-30* mutant showed a modest reduction in the total number of egg + L1 progeny produced compared with WT (Fig. 5B). The number of L4 + adult progeny was also reduced (Fig. 5C). We frequently observed dead L1 larvae on the *hlh-30* mutant plates. Indeed, the percentage of progeny produced (the eggs + L1 count) that survived to the L4 stage was also reduced in the *hlh-30* mutant, though the survival rate varied among individual P0s (Fig. 5D). There was no apparent difference between WT and the *hlh-30* mutant in the timing of egg laying, the number of unfertilized oocytes laid, the total oocyte number (the sum of the egg + L1 and oocytes laid counts), or the rate of embryonic lethality (Fig. 5E, Additional file 7 – Fig. A-C).

Ubiquitous expression of wild-type *hlh-30* rescued both the total brood size and progeny survival through larval development to WT levels (Fig. 5F-H, Additional file 7 – Fig. S4D-E). Together, these results indicate that HLH-30 is required for reproductive fitness in two regards: production of fertilized progeny and survival through larval development.

### Both parental and progeny HLH-30 likely contribute to survival through larval development

The partial larval lethality observed in the *hlh-30* mutant could arise from a requirement for HLH-30 in the parental generation, the developing larvae, or both. To distinguish between these possibilities, we used the Auxin-Inducible Degradation (AID) system to conditionally deplete HLH-30. We inserted the AID degron sequence into the *hlh-30* genomic locus to tag the C-terminus of the HLH-30 protein, allowing its degradation in the presence of auxin and transgenically expressed TIR1 E3 ubiquitin ligase [76]. We treated *hlh-30::aid; eft-3p[ubiquitous]::tir1* animals with auxin in the parental generation (P0), the progeny generation (F1), both, or neither (Fig. 6A). The mean larval survival rate was reduced when HLH-30 was depleted from the parents, the progeny, or both, though the reduction was not statistically significant for the progeny depletion (Fig. 6B). These results suggest that HLH-30 in both the parent and the progeny may contribute to survival through larval development.

**Fig. 5.**
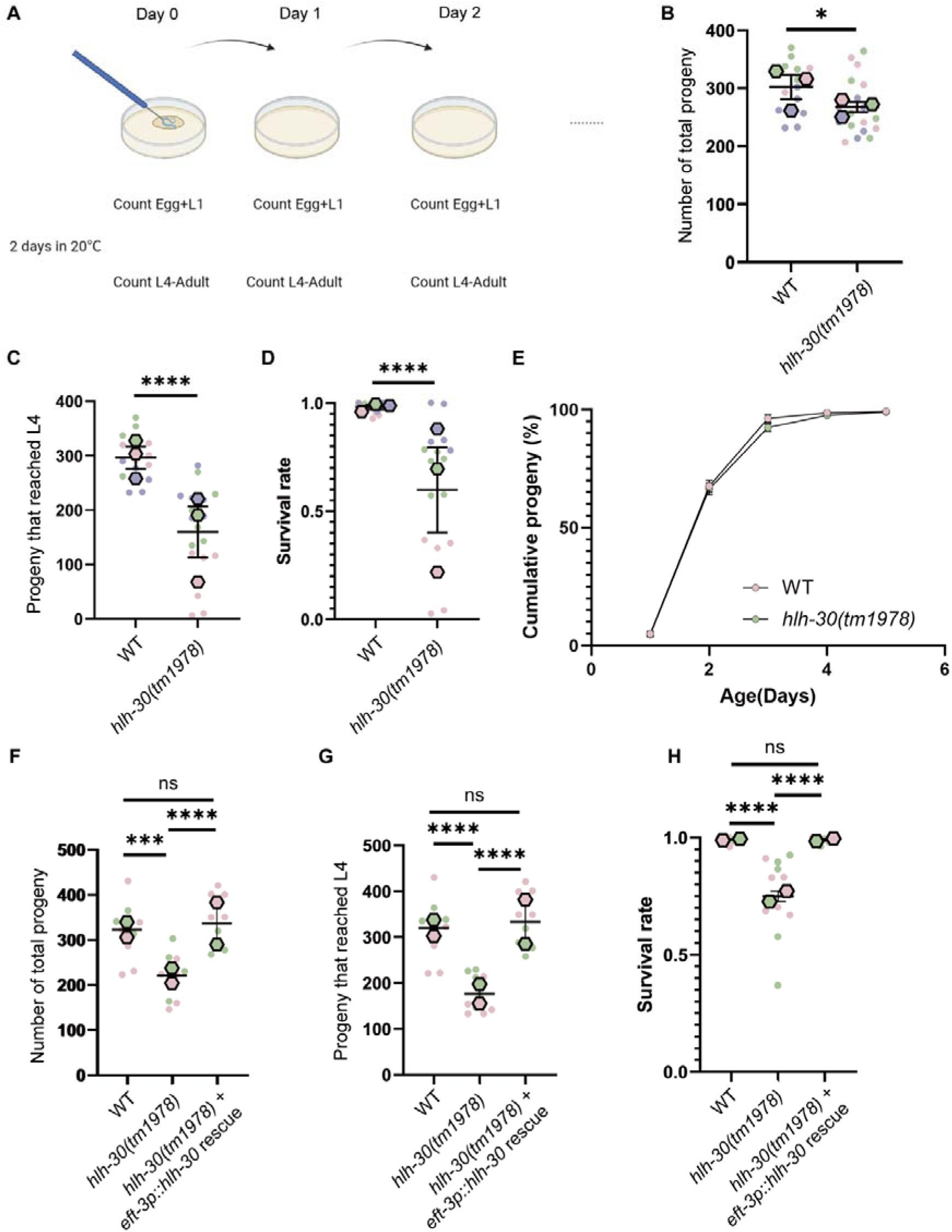
HLH-30 is required for brood size and larval survival. (A) Schematic of the fecundity assay. P0 hermaphrodites were transferred daily starting L4 until reproduction ended. Progeny production was quantified immediately after transfer, and progeny survival was quantified 2 days after transfer. (B-D) The *hlh-30* mutant showed reproductive defects in total number of egg + L1 progeny produced (B), total number of progeny that survived to the L4 stage (C), and progeny survival rate (D) compared to WT. *P<0.05, ****P<0.0001. Student’s *t* test. (E) There was no significant difference in reproductive timing between WT and the *hlh-30* mutant. (F–H) Ubiquitous expression of wild-type *hlh-30* rescued the reproductive defects of the *hlh-30* mutant in total number of egg + L1 progeny produced (F), total number of progeny that survived to L4 (G), and progeny survival rate (H). ***P<0.001, ****P<0.0001, ns: not significant. One-way ANOVA with Tukey post-test. Data were combined across 2-3 independent experimental trials, indicated by the color coding, and the larger hexagon indicates the mean for each trial.

**Fig. 6.**
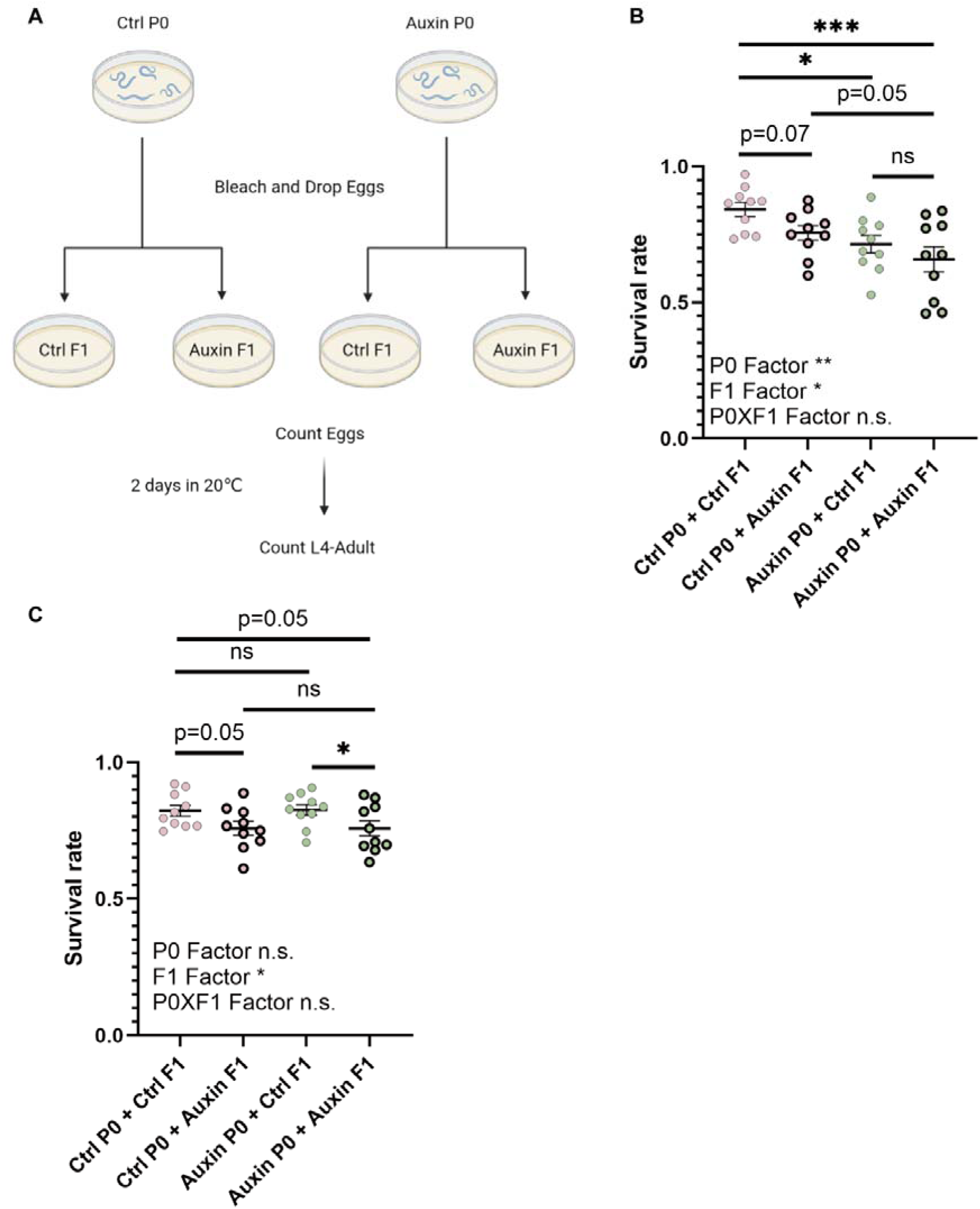
HLH-30 functions in both the parent and progeny, in part from the muscle, to promote progeny survival through larval development. (A) Schematic of the L1 survival assay. P0 worms growing on auxin or control (Ctrl) plates were bleached to extract eggs, which were distributed on auxin or control plates. The number of eggs was counted, and the number of animals that survived to L4 was counted 2 days later. (B) Ubiquitous depletion of HLH-30 in either parental worms or F1 progeny reduced the mean larval survival rate. *P<0.05, **P<0.01, ***P<0.001, ns: not significant. Two-way ANOVA with Sidak post-test. (C) Muscle-specific depletion of HLH-30 in F1 progeny, but not in parental worms alone, led to reduced larval survival. *P<0.05, ns: not significant. Two-way ANOVA with Sidak post-test.

In a pilot experiment investigating the tissue-specific requirement for HLH-30, we depleted HLH-30::AID in neurons, muscle, hypodermis, intestine, or ubiquitously using tissue-specific expression of TIR1. Muscle-specific depletion of HLH-30 resulted in a modest reduction in progeny survival (Additional file 8 - Supplementary Table 2). We therefore tested parental and progeny requirements using *hlh-30::aid; unc-54p[muscle]::tir1* animals. Muscle-specific HLH-30 depletion in the F1 progeny, but not the parental generation alone, reduced progeny survival (Fig. 6C).

### The *hlh-30* mutant shows reduced sperm number compared to WT

Based on the transcriptomic analyses and fecundity assay, we hypothesized that HLH-30 is required to promote some aspect of germ cell generation. To test this, we performed DAPI staining to observe the hermaphrodite germline in Day 1 WT and *hlh-30* mutant adults (Fig. 7A). The relative abundance of germ cells undergoing mitotic proliferation and meiotic differentiation can be assessed based on their distinctive chromatin morphologies and spatial separation along the gonad, and sperm can be identified by their localization and compact nuclei. We observed no apparent change in the lengths of the mitotic or meiotic regions (Fig. 7B-D). However, the *hlh-30* mutant had a reduced average number of sperm cells per gonad arm compared to WT (Fig. 7E). As the number of sperm is limiting for brood size in self-fertilizing *C. elegans* hermaphrodites [77], this reduction may contribute to the reduced brood size in the *hlh-30* mutant.

**Fig. 7.**
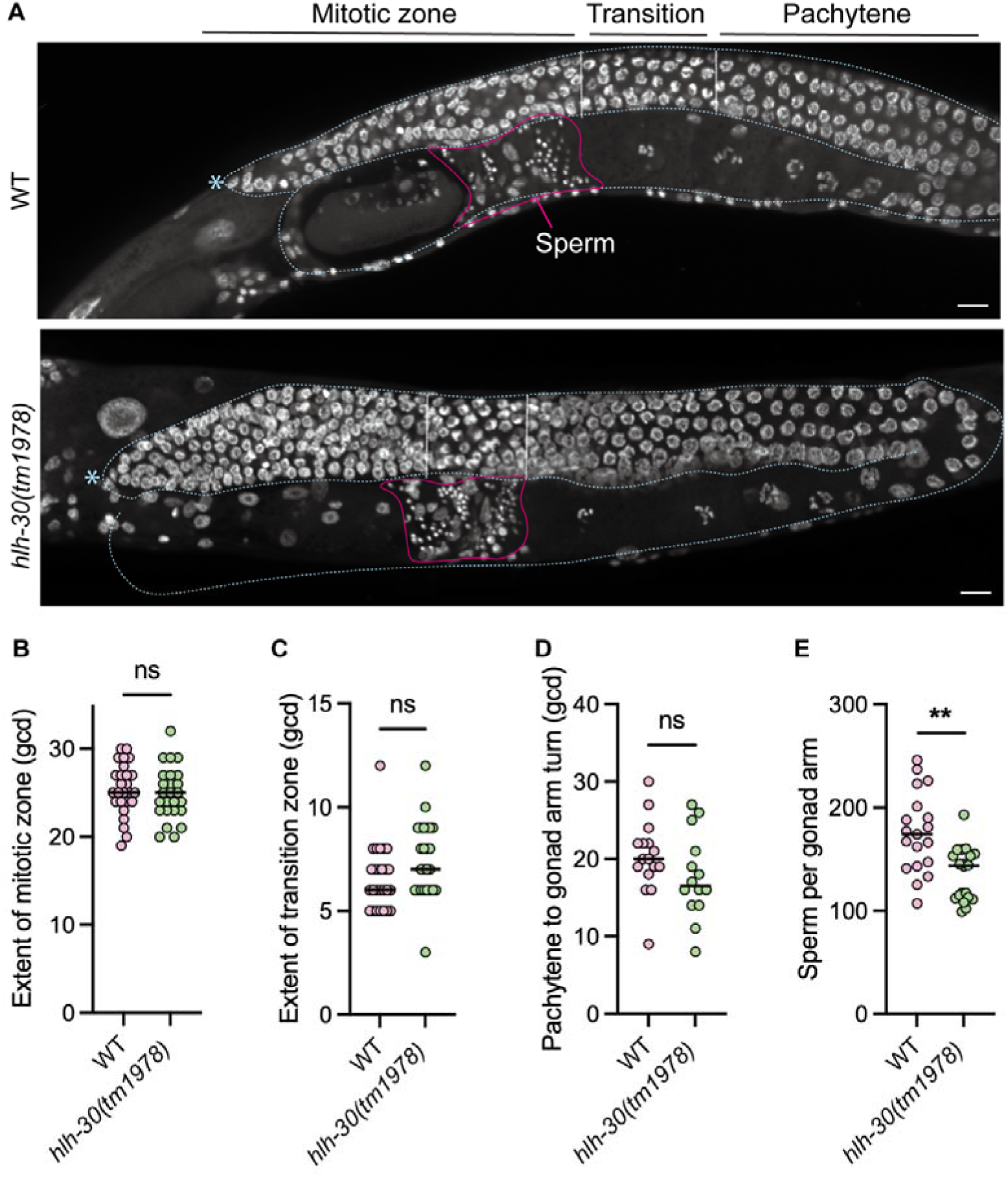
The *hlh-30* mutant shows grossly normal germ cells but reduced sperm number by DAPI staining. (A) Example images of the DAPI-stained germline in Day 1 WT and *hlh-30(tm1978)* mutant adults. Dashed blue line outlines one of a hermaphrodite’s two gonad arms. Asterisk: distal tip cell. Vertical dotted lines mark the beginning of the transition zone, identified by crescent-shaped chromatin, and the boundary between the transition zone and pachytene. Scale bar: 10 μm. (B-D) Quantification of mitotic and meiotic region length, measured in germ cell diameters (gcd). (E) Number of sperm per gonad arm. **P<0.01, ns: not significant; unpaired t-test with Bonferroni correction for the four quantifications shown.

### Elevated *ztf-28* expression contributes to the reproductive defects of *hlh-30* mutant

We next aimed to connect the *hlh-30* mutant’s DEGs to its fitness defects. Among the DEGs identified through the transcriptomic analyses, *ztf-28* was consistently up-regulated across all stages in the *hlh-30* mutant (Fig. 1E), and *ztf-28* has been reported to be expressed predominantly in the germline [78, 79]. We assessed ZTF-28 protein localization and abundance using endogenously tagged ZTF-28::GFP in WT and the *hlh-30* mutant (Fig. 8A)[79]. In WT, we observed ZTF-28::GFP expression predominantly in the germline, where it localized to the germ cell nuclei (Fig. 8A). The ZTF-28::GFP expression pattern and localization were unchanged in the *hlh-30* mutant, but its expression level was higher compared to WT (Fig. 8A-B).

**Fig. 8.**
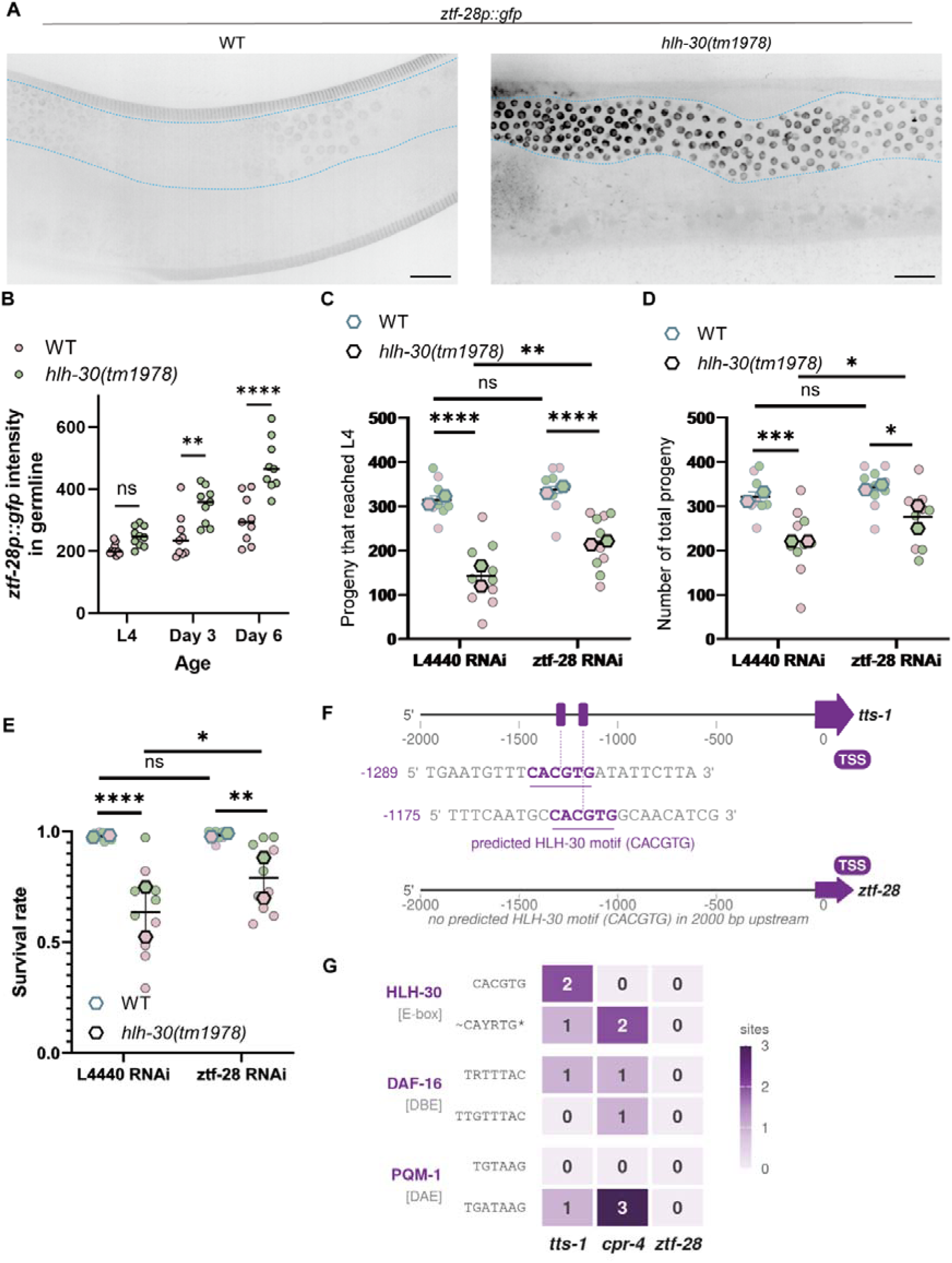
Elevated ZTF-28 expression contributes to the reproductive defects of *hlh-30* mutant animals. (A) Endogenously tagged ZTF-28::GFP showed germline expression and nuclear localization. Scale = 20 μm. (B) ZTF-28::GFP expression was higher in the *hlh-30* mutant compared to WT. (C-E) RNAi-mediated knockdown of *ztf-28* partially rescued the reproductive defects of the *hlh-30* mutant in total number of progeny produced (C), total number of progeny that survived to L4 (D), and progeny survival rate (E). *P<0.05, **P<0.01, ***P<0.001, ****P<0.0001, ns: not significant. Two-way ANOVA with Sidak post-test. Data were combined across 2 independent experimental trials, indicated by the color coding, and the larger hexagon indicates the mean for each trial. (F-G) Analysis of HLH-30 binding sites suggests that *ztf-28* is unlikely to be a direct transcriptional target of HLH-30.

We hypothesized that the aberrant upregulation of *ztf-28* contributes to the reproductive defects caused by loss of HLH-30. Indeed, RNAi-mediated knockdown of *ztf-28* partially rescued the brood size and progeny viability defects in the *hlh-30* mutant, suggesting that elevated *ztf-28* expression contributes to these reproductive defects (Fig. 8C-E).

To assess whether HLH-30 directly regulates *ztf-28* transcription, we examined published HLH-30 ChIP-seq data together with promoter motif analysis. Significant HLH-30 ChIP-seq peaks were detected within the promoter regions of the established HLH-30 targets *tts-1* and *cpr-4,* but not *ztf-28* (Additional file 7 – Fig. S5A-B) [65, 80]. We also searched the promoters of these genes for the HLH-30 E-box and for the DAF-16-binding element (DBE) and DAF-16-associated element (DAE), which are enriched at HLH-30 binding sites [34]. Although multiple such sites were identified within the promoters of *tts-1* and *cpr-4*, none of these sites were detected in the promoter region of *ztf-28* (Fig. 8F-G). These results suggest that HLH-30 may regulate *ztf-28* indirectly.

### HLH-30 regulates autophagy and cellular homeostasis programs during early larval **development**

To investigate the molecular basis of the L1 survival defects in the *hlh-30* mutants, we performed bulk RNA-seq on L1 larvae of WT and the *hlh-30* mutant. PCA and sample correlation analyses showed that L1 samples were separated from L4 and adult samples, reflecting distinct developmental differences in gene expression (Additional file 7 – Fig. S6A-B). PCA restricted to the L1 samples showed that PC1, which accounted for 54.04% of the variance, was associated with loss of *hlh-30* (Fig. 9A). A pairwise comparison between L1 WT versus the *hlh-30* mutant identified 502 DEGs, including 161 up- and 341 downregulated DEGs in the *hlh-30* mutant (Fig. 9B, Additional file 1). Thus, loss of *hlh-30* had a greater transcriptional impact during early larval development than at L4, consistent with its importance for larval survival.

**Fig. 9.**
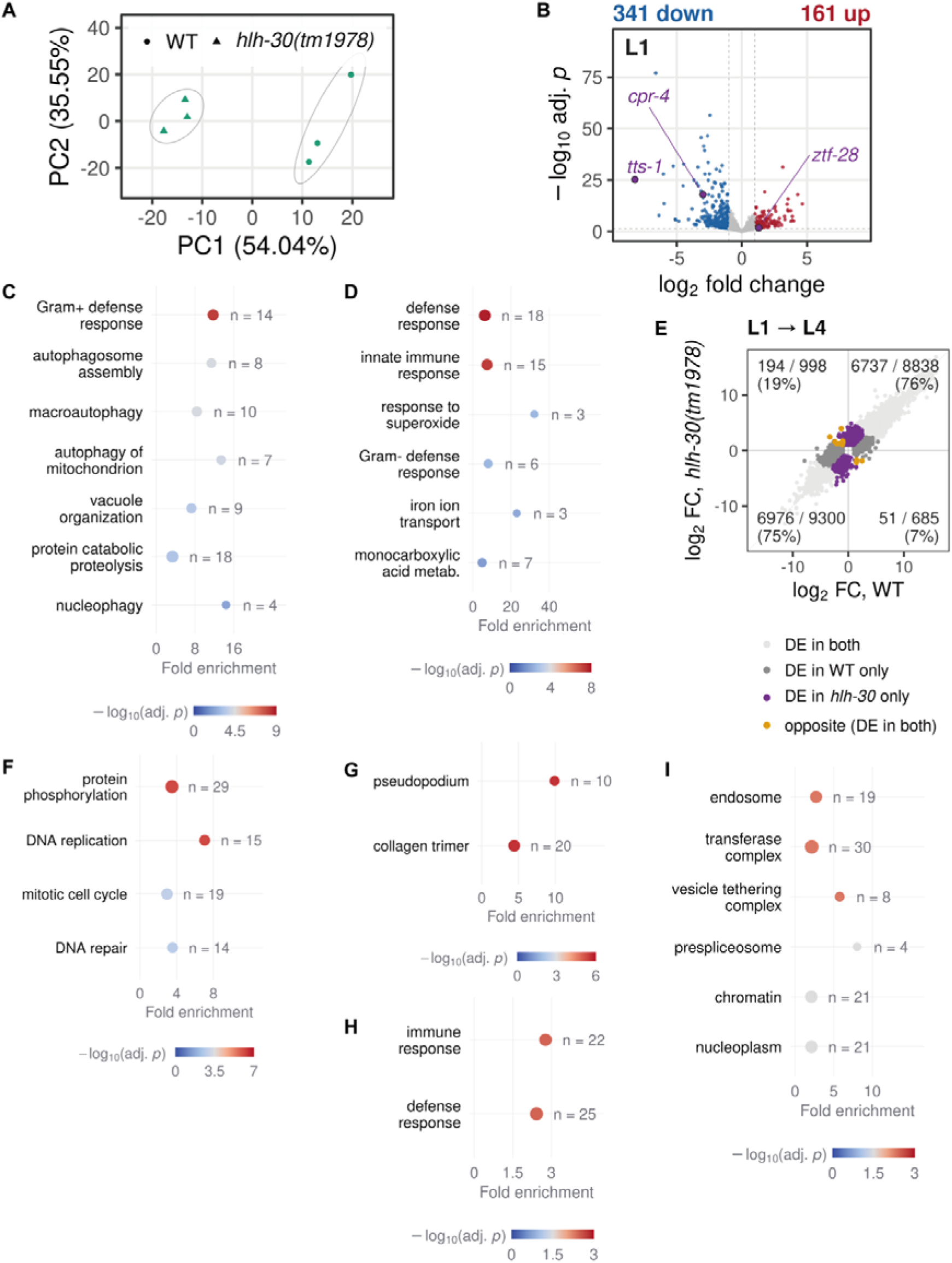
HLH-30 regulates distinct transcriptional programs during larval development. (A) PCA of WT and *hlh-30* mutant RNA-seq samples at L1, which separate by genotype. (B) Volcano plot showing DEGs between WT and the *hlh-30* mutant at L1. (C-D) GO Biological Process enrichment analysis of genes downregulated (C) and upregulated (D) in the *hlh-30* mutant compared to WT at L1. (E) Comparison of DEGs during the L1-to-L4 transition in WT and the *hlh-30* mutant. (F–G) GO enrichment analysis of genes that were upregulated from L1 to L4 more in WT than in the *hlh-30* mutant, or upregulated in WT and downregulated in the mutant. Enriched Biological Process (F) and Cellular Component (G) terms are shown. (H) GO Biological Process enrichment of genes exclusively downregulated during the L1-to-L4 transition in the *hlh-30* mutant. (I) GO Cellular Component enrichment of genes exclusively upregulated during the L1-to-L4 transition in WT.

GO enrichment analysis revealed that genes with decreased expression in the *hlh-30* mutant at L1 were enriched for autophagy- and lysosome-related processes (Fig. 9C). Notably, the role for basal HLH-30 in promoting expression of autophagosome and autophagy initiation genes appeared to be specific to the L1 stage among the four ages examined in this study, and only a small subset of lysosomal protease (cathepsin) genes were dependent on *hlh-30* in adulthood (Additional file 7 – Fig. S7-8, Additional file 5). Genes upregulated in the *hlh-30* mutant at L1 were enriched for defense and innate immune responses, including responses to bacteria and superoxide (Fig. 9D). With our protocol to generate just-hatched, synchronized L1s, a subset of the L1 larvae would have been starved for a maximum of 5 hours. We therefore also analyzed the WT vs *hlh-30(tm1978)* mutant L1 RNA-seq dataset from Murphy et al., in which the L1s were not tightly synchronized but were not starved. This independent dataset showed similar GO enrichments, including autophagy-related processes among genes downregulated in the *hlh-30* mutant (Additional file 4)[25].

To further characterize transcriptional dynamics across development, we performed k-means clustering across all four stages. We identified 6 major expression profiles, most of which were predominantly characterized by developmental changes shared between WT and the *hlh-30* mutant (Additional file 7 – Fig. S6C-D, Additional file 2).

Comparative differential expression analysis identified 9,392 genes that changed by a similar amount during the L1-to-L4 transition in both genotypes, comprising 73% and 77% of the total differentially expressed genes in WT and *hlh-30* mutants, respectively (Fig. 9E, Additional file 3). These analyses suggest that the overall L1-to-L4 developmental transcriptional transition is largely preserved in the *hlh-30* mutant.

GO enrichment analysis of genes that either showed greater upregulation in WT or were upregulated in WT but downregulated in the *hlh-30* mutant revealed enrichment for DNA replication, protein phosphorylation, DNA repair, and the mitotic cell cycle (Fig. 9F), as well as the cellular component pseudopodium (Fig. 9G). These findings suggest that loss of HLH-30 may affect germ cell development. Genes downregulated from L1 to L4 in the *hlh-30* mutant but not in WT were enriched for defense and immune responses (Fig. 9H). Finally, genes exclusively upregulated from L1 to L4 in WT were enriched for endosome-related terms, consistent with the known role of TFEB/HLH-30 in regulating the endo-lysosomal system (Fig. 9I).

### Evidence for age-dependent differences in direct and indirect transcriptional regulation by HLH-30

To investigate whether the age-dependent transcriptional effects of *hlh-30* loss reflect direct or indirect regulation, we examined HLH-30 binding motifs and published ChIP-seq data for the genes that showed differential expression at each stage (Additional file 7 – Fig. S5C, S9). First, we scanned the DEGs’ promoters for the HLH-30 E-box motif from JASPAR (MA1449.1, consensus TCACGTGA; Additional file 7 – Fig. S5C). Second, we examined published ChIP-seq data for evidence of HLH-30 binding at the promoter regions of these genes (Additional file 7 – Fig. S9A). Both approaches indicated that genes with reduced expression in the *hlh-30* mutant (down-DEGs) at L1 were most enriched for evidence of direct HLH-30 regulation (Additional file 7 – Fig. S5C, S9A). Down-DEGs at Day 3 were also enriched for HLH-30 binding in the ChIP-seq analysis (Additional file 7 – Fig. S9A), but the motif enrichment at L4 and Day 3 was not significant (Additional file 7 – Fig. S5C). However, the approaches differed in whether Day 6 down-DEGs were enriched for potential HLH-30 binding. In the ChIP-seq analysis, the fraction of down-DEGs with HLH-30 binding at Day 6 was reduced compared to L1 and Day 3 (Additional file 7 – Fig. S9B). In contrast to the down-DEGs, genes with increased expression in the *hlh-30* mutant (up-DEGs) showed no enrichment for evidence of HLH-30 binding at any age and were depleted of HLH-30 binding at L1 and Day 3 in the ChIP-seq analysis (Additional file 7 – Fig. S5C, S9A). These results suggest that direct transcriptional activation by HLH-30 contributes to basal gene expression across ages, particularly at L1, whereas the increased expression of genes in the *hlh-30* mutant is likely to reflect indirect effects.

We also assessed transcription factor binding motif enrichment among down- and up-DEGs at each age using Simple Enrichment Analysis (SEA, MEME Suite) [60, 61] with nematode motifs from JASPAR 2022 [63] and *C. elegans* motifs from CIS-BP [64] (Additional file 7 - Fig. S10, Additional file 6). This analysis identified 20 candidate transcription factors that may contribute to the transcriptional changes in the *hlh-30* mutant, in addition to E-box binding transcription factors (Additional file 7 - Fig. S10). For instance, motifs associated with NHR-25 were enriched among up-DEGs at Day 3 and Day 6, but not at the larval stages, and several other transcription factor motifs showed stage-specific enrichment as well (Additional file 7 - Fig. S10).

Finally, to compare *hlh-30-*dependent gene regulation across experimental manipulations and laboratories, we examined the overlap between DEGs identified in this study and *hlh-30-*dependent DEGs reported in published studies (Additional file 7 - Fig. S11). Our DEGs overlapped with published *hlh-30*-dependent gene lists more than expected by chance, mostly in the same direction and mostly among genes with reduced expression in the *hlh-30* mutant (Additional file 7 - Fig. S11).

## Discussion

Here, we defined the impact of TFEB/HLH-30 on the *C. elegans* transcriptome across four ages under basal, unstressed laboratory growth conditions. Our results suggest that HLH-30 has a relatively modest impact under these conditions and regulates several physiological processes in an age-dependent manner (Fig. 10A). Further, we identified roles for TFEB/HLH-30 in promoting brood size and progeny survival. Our results support a model in which HLH-30 promotes the production of viable progeny through functions in both the parental and progeny generations (Fig. 10B). We also identified *ztf-28* as a potential downstream effector of HLH-30, whose increased expression in the *hlh-30* mutant may contribute to defects in reproductive fitness. Together, these findings extend the known physiological functions of TFEB/HLH-30 beyond its established roles in stress responses and autophagic and lysosomal biogenesis.

**Fig. 10.**
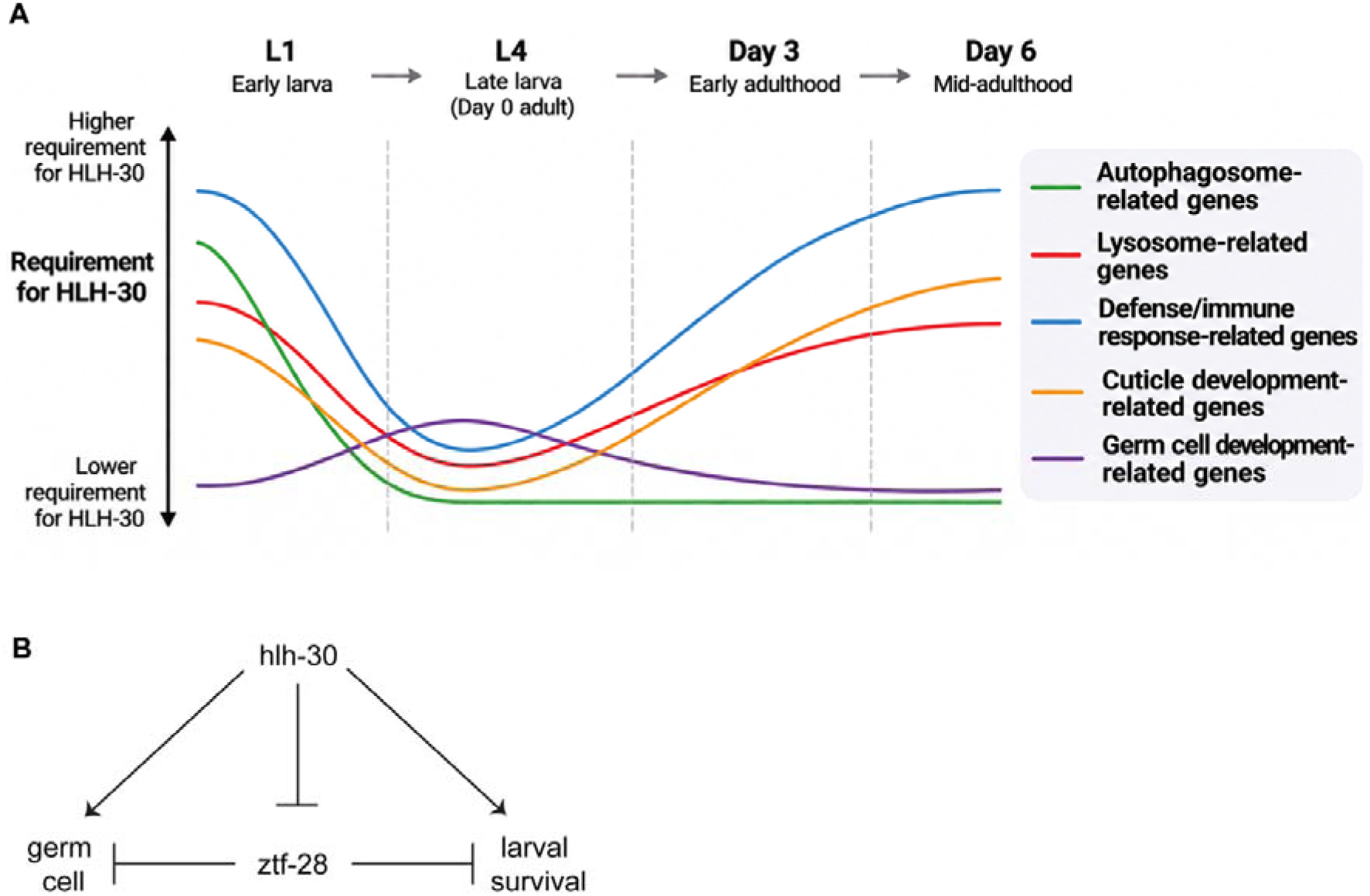
Model for stage-specific functions of HLH-30. (A) The impact of HLH-30 on the transcriptome under unstressed conditions varies with age. (B) HLH-30 promotes reproductive fitness and progeny survival.

Our transcriptomic analyses revealed that the impact of HLH-30 on whole-animal gene expression under unstressed laboratory conditions varies with age in both magnitude and content. Regarding magnitude, the *hlh-30* mutant showed relatively few DEGs at each age, with the greatest number at L1 (Fig. 9B), the least at L4, and increasing numbers from early- to mid-adulthood (Fig. 1E), suggesting that the relative contribution of basal HLH-30 activity to gene expression changes during development. Whether this is due to changes in HLH-30 activity itself, the activity of other transcriptional regulators, or chromatin accessibility remains unknown. Regarding content, while HLH-30 is known to promote autophagy gene expression in response to stress, our results suggest that HLH-30 also promotes basal expression of autophagy and autophagosome-related genes during the L1 stage (Fig. 9C, Additional file 4). However, we found no evidence for a comparable role in regulating basal expression of these genes at L4 or during early to mid-adulthood (Fig. 3-4, Additional file 7 – Fig. S7). HLH-30-dependent expression of a small subset of lysosome-associated genes was observed in both L1 larvae and adults (Additional file 7 – Fig. S8). HLH-30 also had a prominent enrichment for regulating defense and immune response genes under basal conditions in L1 and adulthood, consistent with previous studies demonstrating a role for HLH-30 in innate immunity and pathogen resistance (Fig. 3D, 3G-H, 9C)[23, 31, 33]. Additional enriched processes in early- to mid-adulthood included cuticle production and neuropeptide signaling (Fig. 3C, 3G-H).

Our analyses suggest that age-dependent transcriptional effects of HLH-30 involve both direct and indirect regulation (Fig. 8F-G, Additional file 7 – S5, S9). Based on our assessments for HLH-30 binding motif and ChIP-seq enrichment, genes with reduced expression in the *hlh-30* mutant were enriched for evidence of HLH-30 binding, most prominently at L1 (Additional file 7 – Fig. S5, S9). By contrast, these analyses suggest that genes upregulated in the *hlh-30* mutant are more likely to be indirectly regulated. The enrichment of binding motifs for other transcription factors varied by developmental stage, and those analyses identified several candidates for the regulator(s) of *ztf-28* and the other up-DEGs (Additional file 7 - Fig. S10). Together, these findings suggest that the transcriptional output of basal HLH-30 might involve stage-specific interactions with other regulatory programs.

Beyond its transcriptional functions, we identified a requirement for HLH-30 in reproductive fitness and progeny survival under non-starved conditions. The *hlh-30* mutant’s reduced brood size and partial larval lethality were rescued by ubiquitous expression of wild-type *hlh-30* (Fig. 5F-H), and conditional depletion of HLH-30 further supported its contribution to progeny survival (Fig. 6). Of note, progeny production of the *hlh-30(tm1978)* mutant under standard lab growth conditions has been quantified previously: Parra-Millán et al. reported a mean number of progeny that reached the L4 stage for the *hlh-30* mutant that was approximately 10% smaller than that of WT, a difference that was not statistically significant [73]. Our results show an average reduction of approximately 40%, with considerable variability between batches (Fig. 5C, 5G). This discrepancy in the magnitude of the effect of *hlh-30* loss may reflect differences in environmental conditions between the laboratories, in experimental setup, and/or unknown variation in the strain backgrounds. In addition, our results showed that the *hlh-30* mutant had fewer sperm remaining per gonad arm in young adults, which may contribute to its reduced brood size (Fig. 7E). The reduced sperm number could arise from numerous mechanisms, such as changes in the rate of germ cell proliferation or differentiation, or sperm production or maintenance; DAPI staining analyses do not distinguish between these possibilities.

One potential mediator of the reproductive fitness defects in the *hlh-30* mutant is *ztf-28,* which was consistently upregulated in the *hlh-30* mutant. ZTF-28 is a poorly characterized, putative zinc-finger transcription factor that is predominantly expressed in germ cell nuclei. Reducing *ztf-28* expression by RNAi partially rescues both the brood size and progeny survival defects of the *hlh-30* mutant, suggesting that elevated ZTF-28 contributes to these phenotypes. As analysis of published HLH-30 ChIP-seq data and promoter motif did not find evidence of direct regulation of *ztf-28* by HLH-30, we suggest that HLH-30 limits *ztf-28* expression indirectly. Interestingly, a previous study identified *ztf-28* as a gene for which expression increased with age and knockdown improved healthspan [81]. Therefore, excessive *ztf-28* appears to cause detrimental effects in more than one physiological context. Its mechanism of action in reproductive fitness and healthspan remains unknown, though our data suggest that it does not impact reproductive fitness through widespread transcriptional changes. Though there is no clear mammalian homolog of ZTF-28, approximately 40% of the protein is predicted to be intrinsically disordered [82–84] and approximately 34% comprises predicted droplet-promoting regions [85, 86], making a mammalian functional homolog plausible [87]. Finally, we note several technical limitations of this study. Our fecundity and larval survival assays showed inter-experimental variability, and manual counting of eggs and worms involved some error. These challenges may have reduced the ability to detect modest phenotypes, particularly in the preliminary tissue-specific depletion experiments. Our transcriptomic analyses were performed using bulk RNA-seq, so changes in germ cell-associated gene expression could reflect differences in cell abundance or developmental timing rather than altered transcription within individual cells. For feasibility, adult samples for RNA-seq were prepared by growing *C. elegans* in the presence of FuDR to inhibit progeny generation; that treatment may have impacted the transcriptome in one or both genetic backgrounds [88, 89]. In addition, L4 had three biological replicates per genotype, compared with six at Day 3 and five at Day 6, which reduces the power to detect differential expression at L4 and may partly explain the smaller number of DEGs at this stage. However, the L1 comparison, which also had three replicates per genotype, identified 502 DEGs, so the number of replicates alone does not account for the few DEGs at L4. A subset of animals in our synchronized L1 samples experienced up to several hours of starvation, which is sufficient to activate a transcriptional response in which HLH-30 plays a role [22, 24]. Still, our analysis of an independent dataset of fed WT versus *hlh-30(tm1978)* mutant L1 larvae identified similar GO enrichments to our datasets, including autophagy, which increases confidence that basal expression of these genes indeed depends on *hlh-30* at the L1 stage [25].

## Conclusions

Our work reveals roles for HLH-30 in reproductive fitness and progeny survival under unstressed conditions, extending its physiological functions beyond its known roles in stress responses. Transcriptomic analyses identified age-dependent effects of HLH-30 on several biological processes, including autophagy, lysosomal function, innate immunity, and cuticle development, likely via both direct and indirect transcriptional regulation. We further identified elevated *ztf-28* expression as a contributor to the reproductive defects caused by loss of HLH-30. Together, these findings broaden our understanding of TFEB/HLH-30 as a regulator of organismal physiology and provide a foundation for investigating how its diverse transcriptional functions contribute to development, reproduction, and tissue homeostasis under unstressed conditions.

## Availability of data and materials

The RNA-seq data generated in this study (raw reads and gene counts) are available in the NCBI Gene Expression Omnibus (GEO) under accession GSE354145. Analysis code is available at https://github.com/sherlynpw/hlh30-rnaseq and is archived on Zenodo at https://doi.org/10.5281/zenodo.24213667. All parameters, software versions and quality control steps are given in Additional file 9 - Supplementary Methods.

## Research Funding

Research reported in this publication was supported by the National Institute of General Medical Sciences of the National Institutes of Health under award number R35GM154869, and by the American Heart Association under Award Number 25PRE1366215 to S.P.W. A total of $1,126,290, or 94% percent, of this project was financed with federal funds. A total of 69,548, or 6% percent, of this project was financed with nongovernmental funds. The funders had no role in study design, data collection and analysis, decision to publish, or preparation of the manuscript. The content is solely the responsibility of the authors and does not necessarily represent the official views of the National Institutes of Health.

## Acknowledgments

Some *C. elegans* strains were provided by the *Caenorhabditis* Genetics Center (CGC), which is supported by the National Institutes of Health – Office of Research Infrastructure Program (P40 OD010440). We acknowledge the ENCODE Consortium and the laboratory of Valerie Reinke (Yale University) for generating and making publicly available the ZTF-28::GFP localization images (ENCODE accession ENCSR310EQN), as well as the *ztf-28(st12281[gfp])* allele. We thank Manuel Alvarez for generating the outcrossed *hlh-30(tm1978)* mutant strain.

## Additional Files

Additional file 1 – Differential expression analysis by gene of *hlh-30(tm1978)* versus WT at L1, L4, Day 3, and Day 6 (.xlsx).

Additional file 2 – Genes of each k-means cluster (.xlsx).

Additional file 3 – Comparative differential expression gene lists (.xlsx).

Additional file 4 – GO analysis of the Murphy et al (2019) L1 WT versus *hlh-30(tm1978)*

RNA-seq data (.xlsx).

Additional file 5 – Studies that link each autophagy and lysosome genes to *hlh-30* (.xlsx).

Additional file 6 – All enriched SEA motifs (.xlsx).

Additional file 7 – Supplemental figures. A pdf (.pdf) file with Supplemental Figures 1-11 and their captions.

Additional file 8 – Supplemental Tables 1-2 (.xlsx).

Additional file 9 – Supplementary Methods (.pdf).

