## Additional file 7 - Supp figs for "*Caenorhabditis elegans* HLH-30/TFEB has age-dependent effects on the transcriptome and is required for normal progeny number and larval survival under basal growth conditions"

### Supplemental Figures

**Fig. S1. Additional differential expression and clustering analyses for WT versus *hlh-30(tm1978)* from the L4, Day 3, and Day 6 datasets.** (A) DEGs shared at Day 3 and Day 6 but not the L4 stage. (B) HDBSCAN clustering of 5,044 DEGs among L4, Day 3, and Day 6 WT and *hlh-30* mutant samples identified seven distinct temporal expression profiles and left 523 genes unassigned (noise). (C) PCA of the Day 3 and Day 6 samples as in Fig. 1D, colored by cohort (20 °C or 22 °C).

**S1**

**A**

up in *hlh-30(tm1978)* (13)

| | L4 | D3 | D6 |
|---|---|---|---|
| *C17B7.5* | 0.1 | 2.4 | 1.8 |
| *clec-4* | 0.0 | 1.9 | 2.4 |
| *C04E12.2* | 0.9 | 1.5 | 1.1 |
| *spp-27* | 0.1 | 2.3 | 1.1 |
| *T16H12.2* | 0.3 | 2.0 | 1.1 |
| *C45B2.8* | 0.0 | 1.8 | 1.4 |
| *dyf-3* | 0.2 | 1.1 | 1.4 |
| *clec-66* | 0.0 | 1.3 | 1.2 |
| *C09G5.7* | 0.0 | 1.2 | 1.1 |
| *C14C6.5* | 0.0 | 1.0 | 1.3 |
| *F54E2.1* | 0.2 | 1.1 | 1.0 |
| *C04E12.5* | 0.0 | 1.2 | 1.1 |
| *F25A2.1* | 0.1 | 1.2 | 1.0 |

down in *hlh-30(tm1978)* (34)

| | L4 | D3 | D6 |
|---|---|---|---|
| *ilys-3* | -0.1 | -3.7 | -4.5 |
| *ilys-2* | -0.1 | -1.7 | -5.0 |
| *C08E3.13* | 0.0 | -2.9 | -3.6 |
| *Y53F4B.45* | -0.8 | -2.4 | -3.0 |
| *C08E3.1* | 0.0 | -2.0 | -3.8 |
| *T23E7.6* | 0.0 | -2.2 | -3.4 |
| *linc-17* | -0.3 | -2.8 | -2.4 |
| *F46A8.13* | -0.5 | -2.6 | -2.2 |
| *cpr-4* | -0.2 | -2.0 | -2.7 |
| *C37A5.3* | -0.8 | -1.7 | -2.2 |
| *cyp-13A5* | -0.1 | -1.7 | -2.8 |
| *W09G12.7* | -0.1 | -1.2 | -3.2 |
| *fipr-1* | 0.0 | -1.7 | -2.7 |
| *F55H12.2* | -0.3 | -1.6 | -2.3 |
| *lys-4* | -0.1 | -1.9 | -2.2 |
| *Y105C5B.5* | -0.9 | -1.2 | -2.0 |
| *hlh-30* | -1.0 | -1.3 | -1.6 |
| *cpr-1* | -0.3 | -1.5 | -1.8 |
| *spp-2* | -0.5 | -1.3 | -1.5 |
| *fipr-24* | 0.1 | -1.8 | -1.5 |
| *cyp-34A9* | 0.0 | -1.0 | -2.2 |
| *E02H4.7* | -0.2 | -1.1 | -1.8 |
| *nssp-49* | 0.0 | -1.5 | -1.6 |
| *lys-5* | -0.1 | -1.1 | -1.8 |
| *Y54F10BM.22* | -0.5 | -1.1 | -1.4 |
| *T03F1.11* | 0.1 | -1.2 | -1.8 |
| *fipr-2* | 0.0 | -1.1 | -1.8 |
| *Y37F4.1* | 0.0 | -1.7 | -1.1 |
| *T05E12.3* | -0.1 | -1.1 | -1.6 |
| *T16G1.7* | 0.0 | -1.4 | -1.3 |
| *C37A5.5* | 0.1 | -1.4 | -1.3 |
| *his-61* | -0.1 | -1.1 | -1.0 |
| *clec-218* | 0.0 | -1.1 | -1.1 |
| *his-25* | 0.0 | -1.0 | -1.0 |

log2FC, *hlh-30(tm1978)* vs WT
pale = not a DEG at that stage

**B**

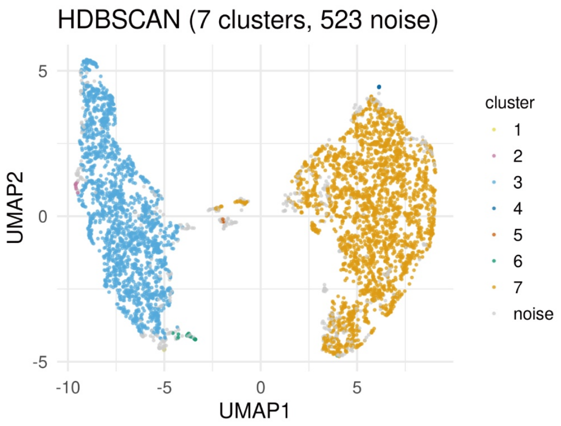


**C**

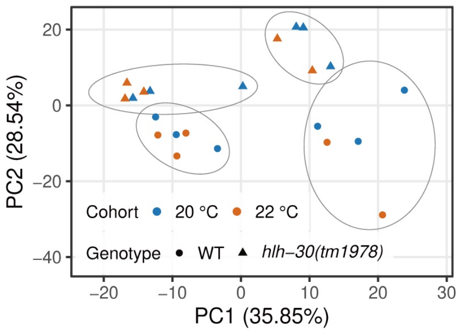

**Fig. S2. Additional enrichment analyses of age-dependent gene expression changes between WT and the *hlh-30(tm1978)* mutant.** (A-B) GO biological process enrichment analysis of genes that were exclusively up- (A) and down-regulated (B) in WT during the L4-to-Day 3 transition. (C-D) GO biological process enrichment analysis of genes that were exclusively down-regulated in the *hlh-30* mutant (C) and up-regulated in WT (D) during the L4-to-Day 6 transition. (E) GO biological process enrichment analysis of genes that were exclusively down-regulated in WT during the Day 3-to-Day 6 transition.

**S2**

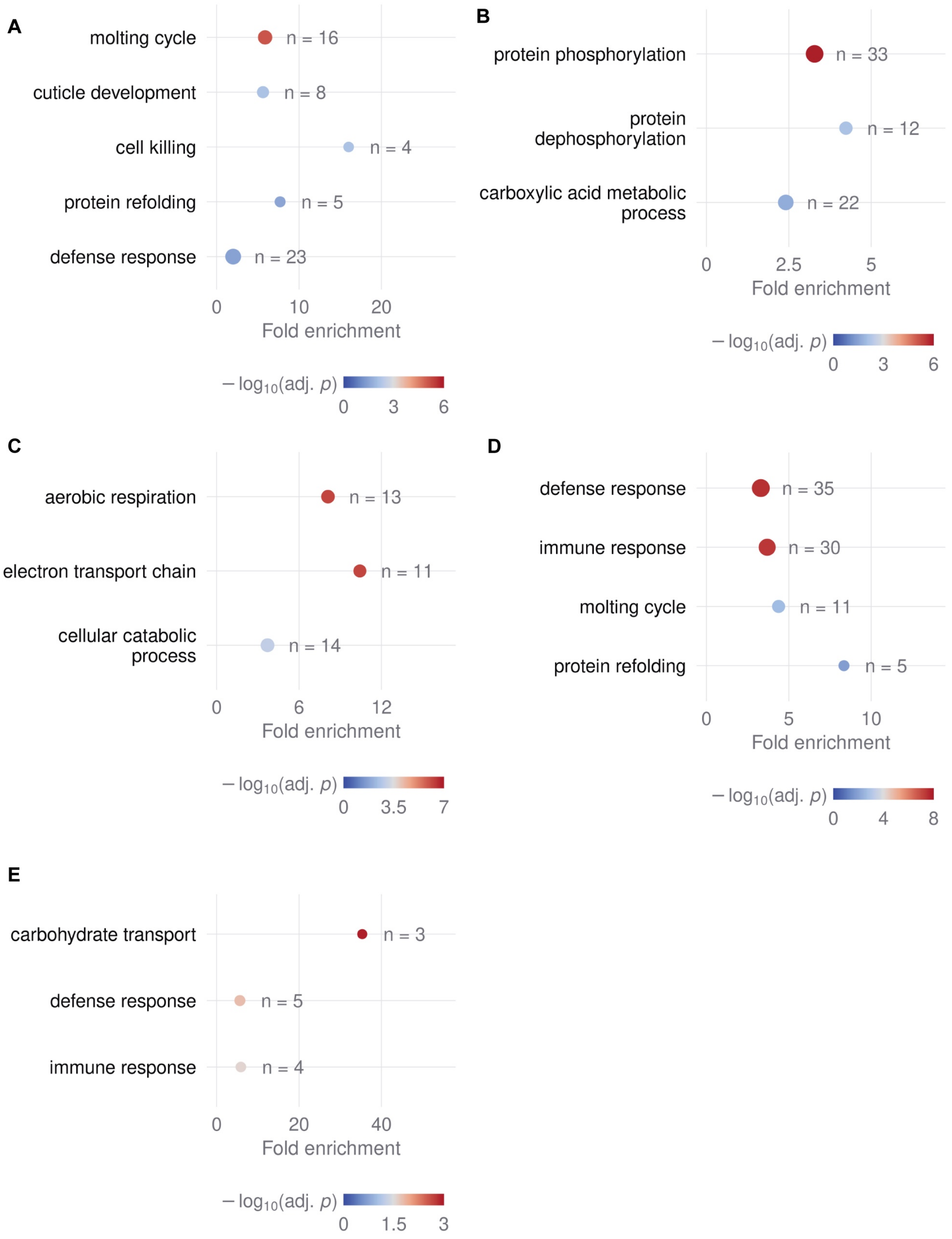
A
molting cycle
n = 16
cuticle development
n = 8
cell killing
n = 4
protein refolding
n = 5
defense response
n = 23
0
10
20
Fold enrichment
− log10(adj. p)
0
3
6
B
protein phosphorylation
n = 33
protein dephosphorylation
n = 12
carboxylic acid metabolic process
n = 22
0
2.5
5
Fold enrichment
− log10(adj. p)
0
3
6
C
aerobic respiration
n = 13
electron transport chain
n = 11
cellular catabolic process
n = 14
0
6
12
Fold enrichment
− log10(adj. p)
0
3.5
7
D
defense response
n = 35
immune response
n = 30
molting cycle
n = 11
protein refolding
n = 5
0
5
10
Fold enrichment
− log10(adj. p)
0
4
8
E
carbohydrate transport
n = 3
defense response
n = 5
immune response
n = 4
0
20
40
Fold enrichment
− log10(adj. p)
0
1.5
3

**Fig. S3. Germline genes trended toward lower expression in the *hlh-30* mutant at L4.** (A-B) Expression of the 31 oocyte and germline genes (A) and the 18 sperm genes (B) in WT and *hlh-30(tm1978)* at L1, L4, Day 3 (D3) and Day 6 (D6). Genes were taken from germ cell-related GO terms enriched in the comparative differential expression gene lists of the adult transitions (L4-to-Day 3, L4-to-Day 6 and Day 3-to-Day 6). The oocyte and germline set holds the genes of terms such as germ cell development, gamete generation and oogenesis, and the sperm set holds the genes of the pseudopodium term, which are mainly major sperm protein genes. Color shows the z-score of each gene's mean expression in each genotype and stage (DESeq2 variance-stabilized counts with replicate and unwanted-variation effects removed; see Methods). Rows are hierarchically clustered. (C-D) Mean z-scored expression of the oocyte and germline genes (C) and the sperm genes (D). Lines show the mean across genes and shaded bands show the SEM. In (D), the band is narrower than the line (SEM 0.01 to 0.02).

S3

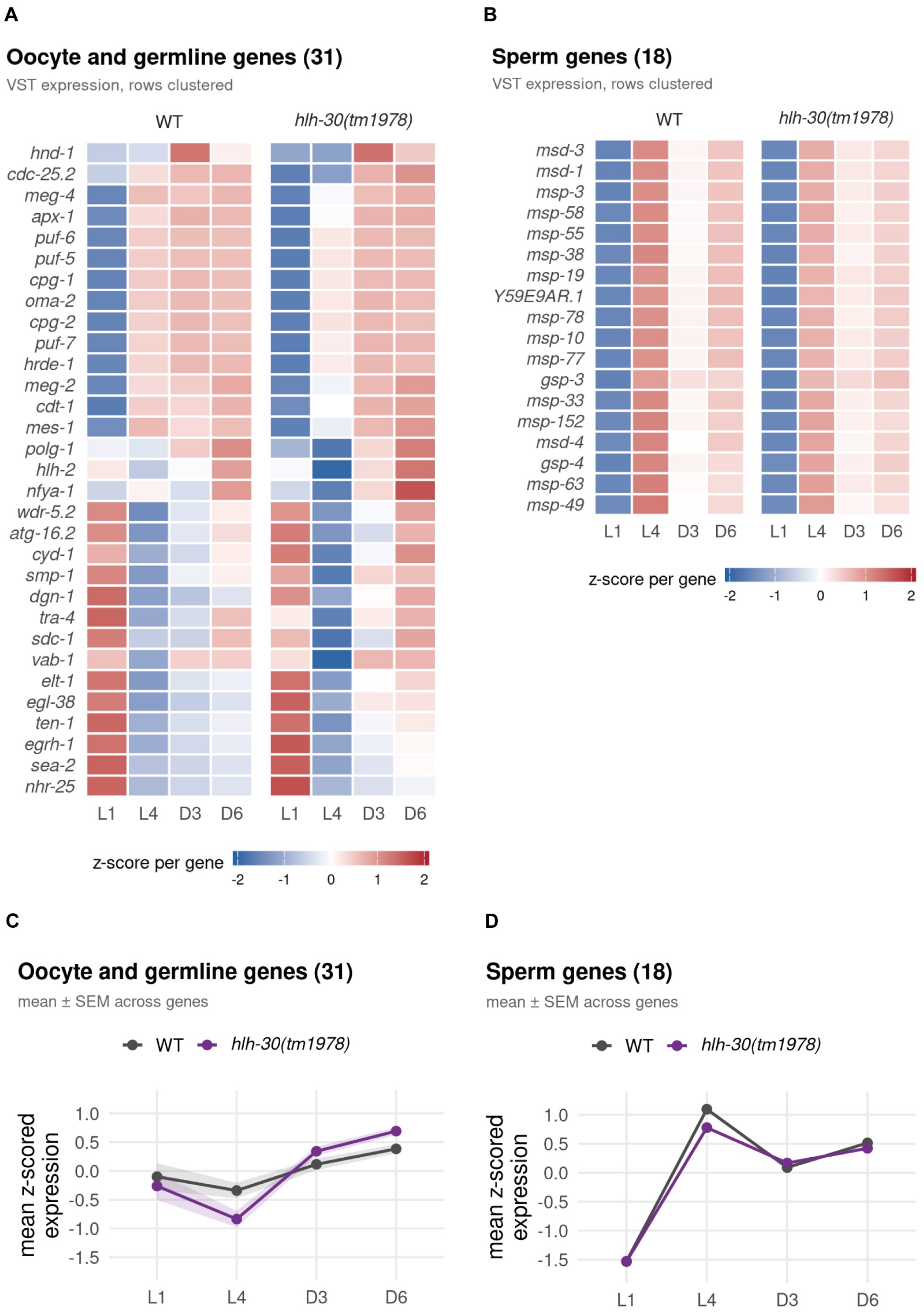
A
Oocyte and germline genes (31)
VST expression, rows clustered
WT
hlh-30(tm1978)
hnd-1
cdc-25.2
meg-4
apx-1
puf-6
puf-5
cpg-1
oma-2
cpg-2
puf-7
hrde-1
meg-2
cdt-1
mes-1
polg-1
hlh-2
nfya-1
wdr-5.2
atg-16.2
cyd-1
smp-1
dgn-1
tra-4
sdc-1
vab-1
elt-1
egl-38
ten-1
egrh-1
sea-2
nhr-25
L1
L4
D3
D6
z-score per gene
-2
-1
0
1
2
B
Sperm genes (18)
VST expression, rows clustered
msd-3
msd-1
msp-3
msp-58
msp-55
msp-38
msp-19
Y59E9AR.1
msp-78
msp-10
msp-77
gsp-3
msp-33
msp-152
msd-4
gsp-4
msp-63
msp-49
C
Oocyte and germline genes (31)
mean ± SEM across genes
mean z-scored expression
1.0
0.5
0.0
-0.5
-1.0
-1.5
D
Sperm genes (18)
mean ± SEM across genes

**Fig. S4. Quantification of additional aspects of progeny production between WT and the *hlh-30(tm1978)* mutant.** (A-C) The *hlh-30* mutant showed no difference in the total number of unfertilized oocytes produced (A), total number of female germ cells (calculated by total number of unfertilized oocytes + total number of progeny produced) (B), and egg hatching rate (C) compared to WT. ns: not significant. Student's *t* test. (D-E) Ubiquitous expression of wild-type *hlh-30* showed no difference in the total number of unfertilized oocytes produced (D), and rescued the reduction in the total number of female germ cells (E) in the *hlh-30* mutant. **P<0.01, ****P<0.0001, ns: not significant. One-way ANOVA with Tukey post-test. Of note, in this trial, the number of total female germ cells was significantly reduced, in contrast to the experiments in S4B. Perhaps this *hlh-30* phenotype is highly variable or varies by subtle environmental conditions in the lab. (F-G) RNAi-mediated knockdown of *ztf-28* had no obvious effect on the total number of unfertilized oocytes produced (F), and the total number of female germ cells (G). *P<0.05, ns: not significant. Two-way ANOVA with Sidak post-test. Data were combined across 2-3 independent experimental trials, indicated by the color coding, and the larger hexagon indicates the mean for each trial.

S4

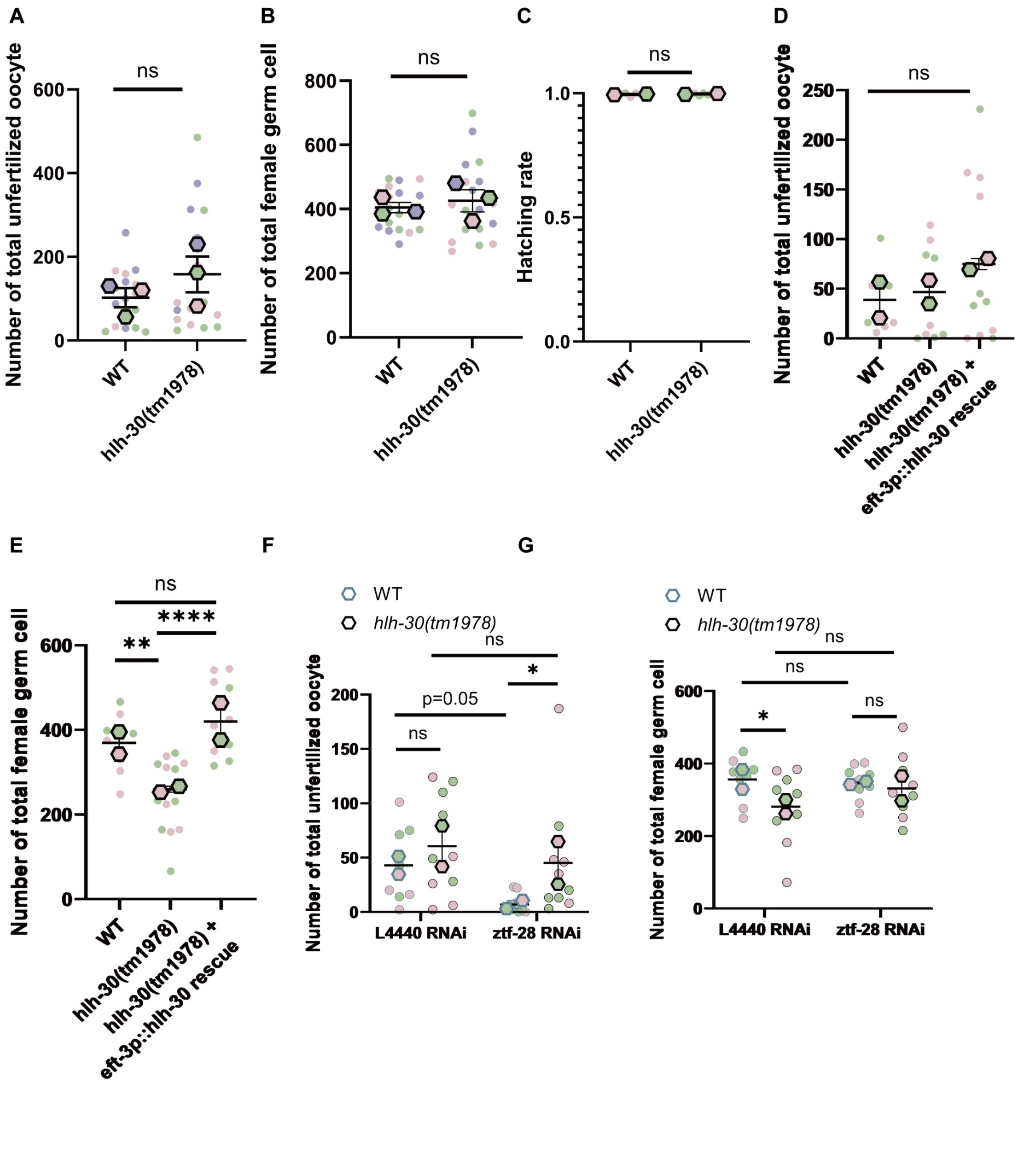
A
ns
Number of total unfertilized oocyte
600
400
200
0
WT
hlh-30(tm1978)
B
ns
Number of total female germ cell
800
600
400
200
0
WT
hlh-30(tm1978)
C
ns
Hatching rate
1.0
0.5
0.0
WT
hlh-30(tm1978)
D
ns
Number of total unfertilized oocyte
250
200
150
100
50
0
WT
hlh-30(tm1978)
hlh-30(tm1978) +
eft-3p::hlh-30 rescue
E
ns
**
****
Number of total female germ cell
600
400
200
0
WT
hlh-30(tm1978)
hlh-30(tm1978) +
eft-3p::hlh-30 rescue
F
WT
hlh-30(tm1978)
ns
*
p=0.05
ns
Number of total unfertilized oocyte
200
150
100
50
0
L4440 RNAi
ztf-28 RNAi
G
WT
hlh-30(tm1978)
ns
ns
*
ns
Number of total female germ cell
600
400
200
0
L4440 RNAi
ztf-28 RNAi

**Fig. S5. HLH-30 binds known targets but not *ztf-28*, and its E-box is enriched in down-regulated but not up-regulated genes.** (A) Mean HLH-30 ChIP-seq signal, as fold enrichment over input (flanks set to 1), within 3 kb of the transcription start sites (TSSs) of all 19,985 protein-coding genes, for two L4 and two late-embryo replicates. (B) HLH-30 ChIP-seq coverage at the *ztf-28*, *tts-1* and *cpr-4* promoters, relative to the genome-wide mean and averaged over the four experiments, on a y-axis shared across loci. Black bars mark MACS2 peaks ($Q < 1 \times 10^{-5}$) in any of the four experiments. Scale bars, 1 kb. ChIP-seq data in (A-B) are from modENCODE [65], obtained from ChIP-Atlas (ce11). (C) Percentage of promoters (−2,000 to +200 bp of the TSS) with a match to the canonical HLH-30 E-box (TCACGTGA, JASPAR MA1449.1, $P < 1 \times 10^{-4}$) among DEGs in *hlh-30(tm1978)* vs WT at each stage (D3, Day 3; D6, Day 6), for all DEGs and separately for up- and down-regulated DEGs. Gray dots and the dashed line mark the background (21.5%), which is the share of all expressed genes with a match. *$q < 0.05$, ***$q < 0.001$, two-sided Fisher's exact test against the background with Benjamini-Hochberg adjustment.

**S5**

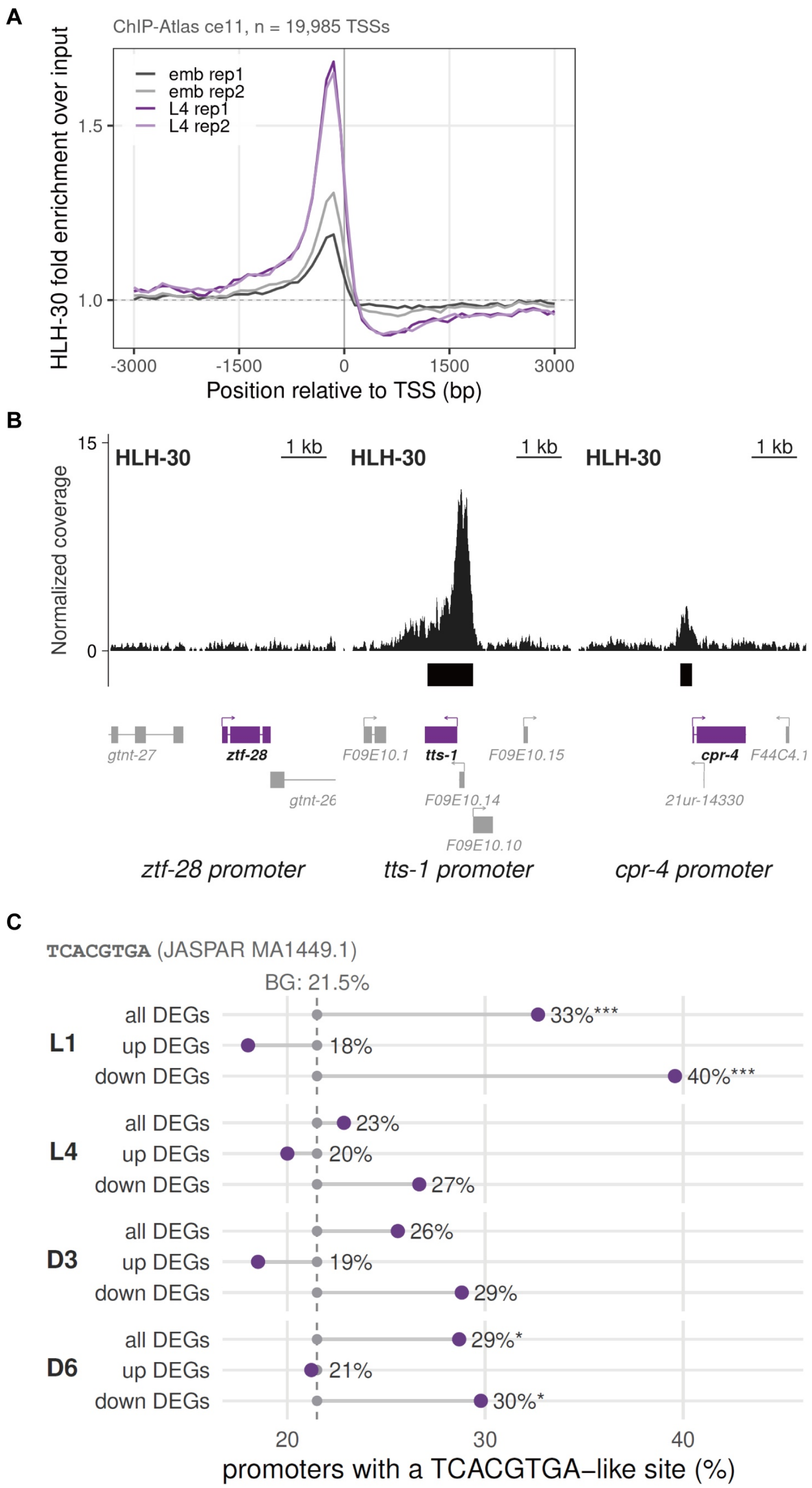
A
ChIP-Atlas ce11, n = 19,985 TSSs
HLH-30 fold enrichment over input
emb rep1
emb rep2
L4 rep1
L4 rep2
1.5
1.0
-3000
-1500
0
1500
3000
Position relative to TSS (bp)
B
15
0
Normalized coverage
HLH-30
1 kb
HLH-30
1 kb
HLH-30
1 kb
gtnt-27
ztf-28
gtnt-26
F09E10.1
tts-1
F09E10.15
F09E10.14
F09E10.10
cpr-4
F44C4.1
21ur-14330
ztf-28 promoter
tts-1 promoter
cpr-4 promoter
C
TCACGTGA (JASPAR MA1449.1)
BG: 21.5%
L1
all DEGs
33%***
up DEGs
18%
down DEGs
40%***
L4
all DEGs
23%
up DEGs
20%
down DEGs
27%
D3
all DEGs
26%
up DEGs
19%
down DEGs
29%
D6
all DEGs
29%*
up DEGs
21%
down DEGs
30%*
20
30
40
promoters with a TCACGTGA–like site (%)

**Fig. S6. Additional clustering and enrichment analyses using the transcriptomic data from WT versus the *hlh-30(tm1978)* mutant at the L1 larval stage.** (A–B) PCA (A) and hierarchical clustering (B) of WT and *hlh-30* mutant RNA-seq samples across developmental and adult stages, highlighting the distinct transcriptional profiles of L1 animals. (C–D) K-means clustering of 16,465 genes across developmental and adult stages in WT and the *hlh-30* mutant identified 6 individual clusters. (E) GO biological process enrichment analysis of cluster 3.

S6

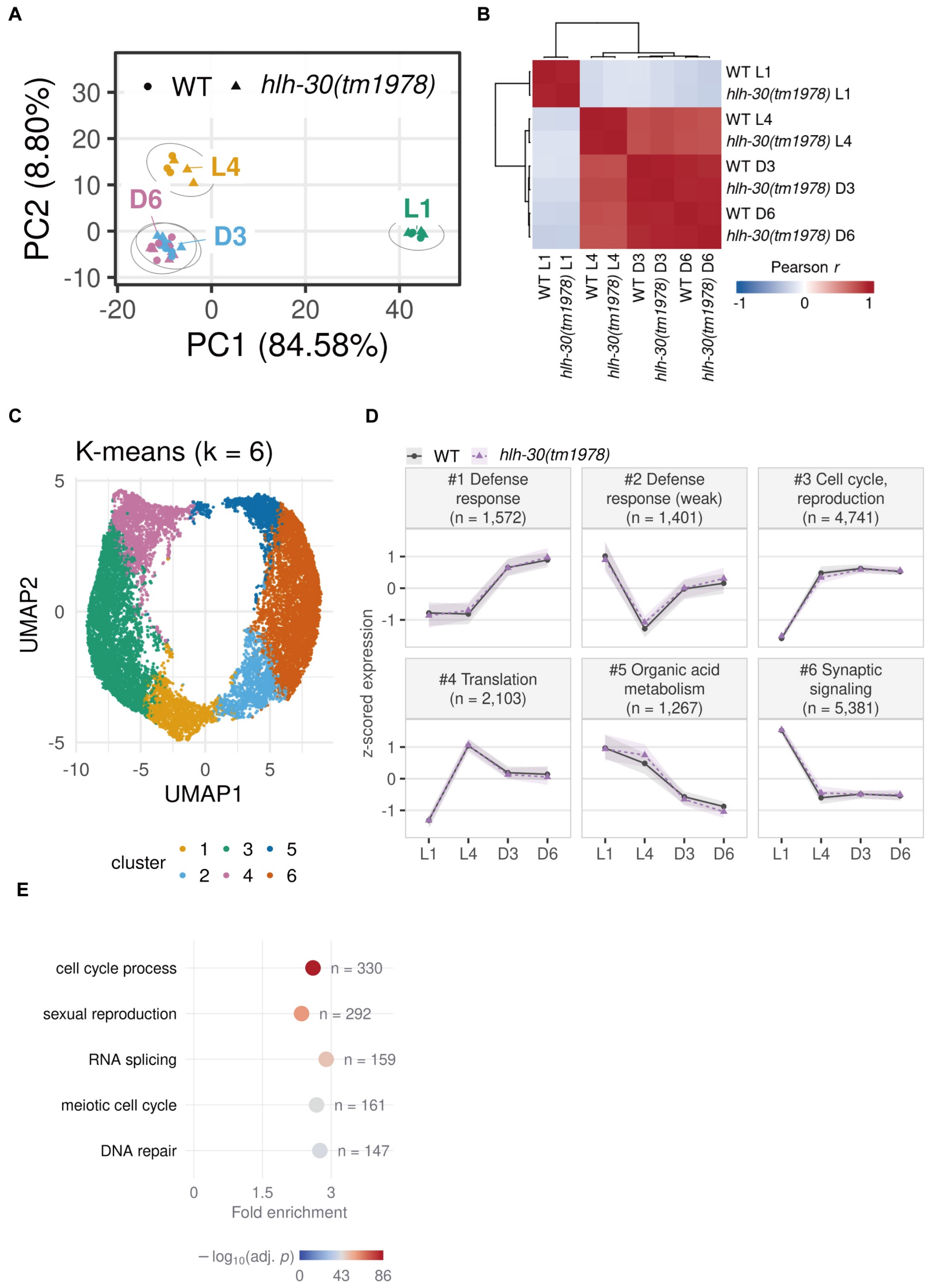
A
WT
hlh-30(tm1978)
PC2 (8.80%)
PC1 (84.58%)
L4
D6
D3
L1
B
WT L1
hlh-30(tm1978) L1
WT L4
hlh-30(tm1978) L4
WT D3
hlh-30(tm1978) D3
WT D6
hlh-30(tm1978) D6
Pearson r
C
K-means (k = 6)
UMAP2
UMAP1
cluster
D
#1 Defense response (n = 1,572)
#2 Defense response (weak) (n = 1,401)
#3 Cell cycle, reproduction (n = 4,741)
#4 Translation (n = 2,103)
#5 Organic acid metabolism (n = 1,267)
#6 Synaptic signaling (n = 5,381)
z-scored expression
L1 L4 D3 D6
E
cell cycle process n = 330
sexual reproduction n = 292
RNA splicing n = 159
meiotic cell cycle n = 161
DNA repair n = 147
Fold enrichment
− log10(adj. p)

**Fig. S7. Core autophagy genes are lower in the *hlh-30* mutant at L1 but not at L4 or in adults.** (A) The 21 core autophagy genes that published studies show to need *hlh-30* for their expression, a subset of (B). The number in parentheses is the number of studies, which are listed in Supplementary File 5. (B) All 69 genes of the *C. elegans* core autophagy gene table of Zhang and Meléndez [67], regardless of *hlh-30*, grouped by role. In (A-B), color shows the $\log_2$ fold change in *hlh-30(tm1978)* vs WT at each stage (D3, Day 3; D6, Day 6). Two asterisks mark DEGs (adjusted $P < 0.05$ and $|\log_2 FC| \geq 1$), and one asterisk marks genes with adjusted $P < 0.05$ but $|\log_2 FC| < 1$. Gray cells mark genes that were not tested at that stage. *hlh-30* itself (labeled “deleted”), which carries the *tm1978* deletion, is a DEG at every stage and serves as a positive control.

S7

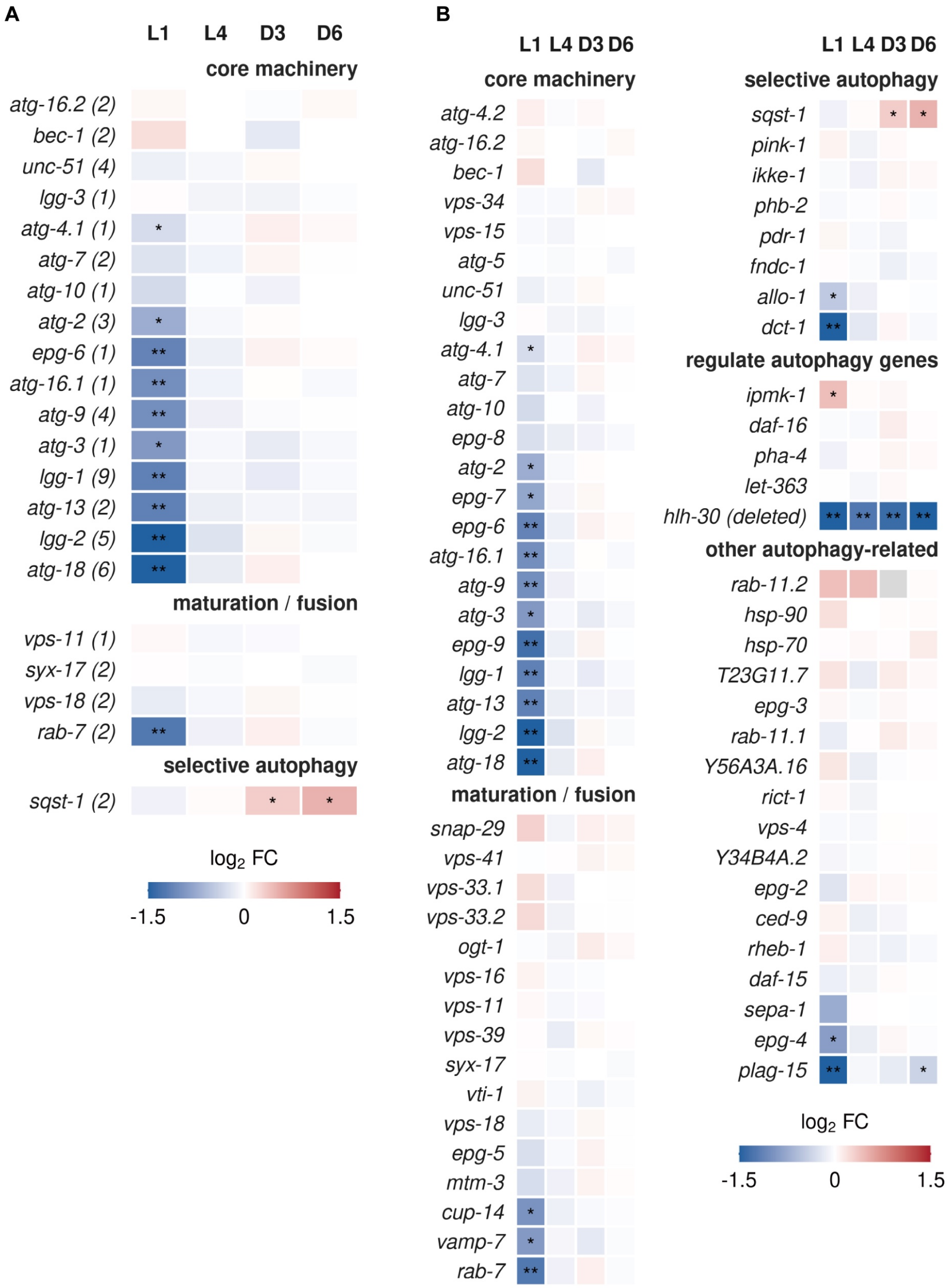

A
L1 L4 D3 D6
core machinery
atg-16.2 (2)
bec-1 (2)
unc-51 (4)
lgg-3 (1)
atg-4.1 (1)
atg-7 (2)
atg-10 (1)
atg-2 (3)
epg-6 (1)
atg-16.1 (1)
atg-9 (4)
atg-3 (1)
lgg-1 (9)
atg-13 (2)
lgg-2 (5)
atg-18 (6)
maturation / fusion
vps-11 (1)
syx-17 (2)
vps-18 (2)
rab-7 (2)
selective autophagy
sqst-1 (2)
log2 FC
-1.5 0 1.5
B
L1 L4 D3 D6
core machinery
atg-4.2
atg-16.2
bec-1
vps-34
vps-15
atg-5
unc-51
lgg-3
atg-4.1
atg-7
atg-10
epg-8
atg-2
epg-7
epg-6
atg-16.1
atg-9
atg-3
epg-9
lgg-1
atg-13
lgg-2
atg-18
maturation / fusion
snap-29
vps-41
vps-33.1
vps-33.2
ogt-1
vps-16
vps-11
vps-39
syx-17
vti-1
vps-18
epg-5
mtm-3
cup-14
vamp-7
rab-7
L1 L4 D3 D6
selective autophagy
sqst-1
pink-1
ikke-1
phb-2
pdr-1
fndc-1
allo-1
dct-1
regulate autophagy genes
ipmk-1
daf-16
pha-4
let-363
hlh-30 (deleted)
other autophagy-related
rab-11.2
hsp-90
hsp-70
T23G11.7
epg-3
rab-11.1
Y56A3A.16
rict-1
vps-4
Y34B4A.2
epg-2
ced-9
rheb-1
daf-15
sepa-1
epg-4
plag-15
log2 FC
-1.5 0 1.5

**Fig. S8. Cathepsin genes are lower in the *hlh-30* mutant at L1, Day 3 and Day 6.** (A) The 27 lysosome genes that published studies show to need *hlh-30* for their expression, a subset of (B). The number in parentheses is the number of studies, which are listed in Supplementary File 5. (B) All 85 lysosome genes of Sun et al. [68], in their five classes: membrane proteins, V-ATPase V0 subunits, V-ATPase V1 subunits, cathepsins and other hydrolases. In (A-B), color shows the $\log_2$ fold change in *hlh-30(tm1978)* vs WT at each stage (D3, Day 3; D6, Day 6). Two asterisks mark DEGs (adjusted $P < 0.05$ and $|\log_2 FC| \geq 1$), and one asterisk marks genes with adjusted $P < 0.05$ but $|\log_2 FC| < 1$. Gray cells mark genes that were not tested at that stage.

S8

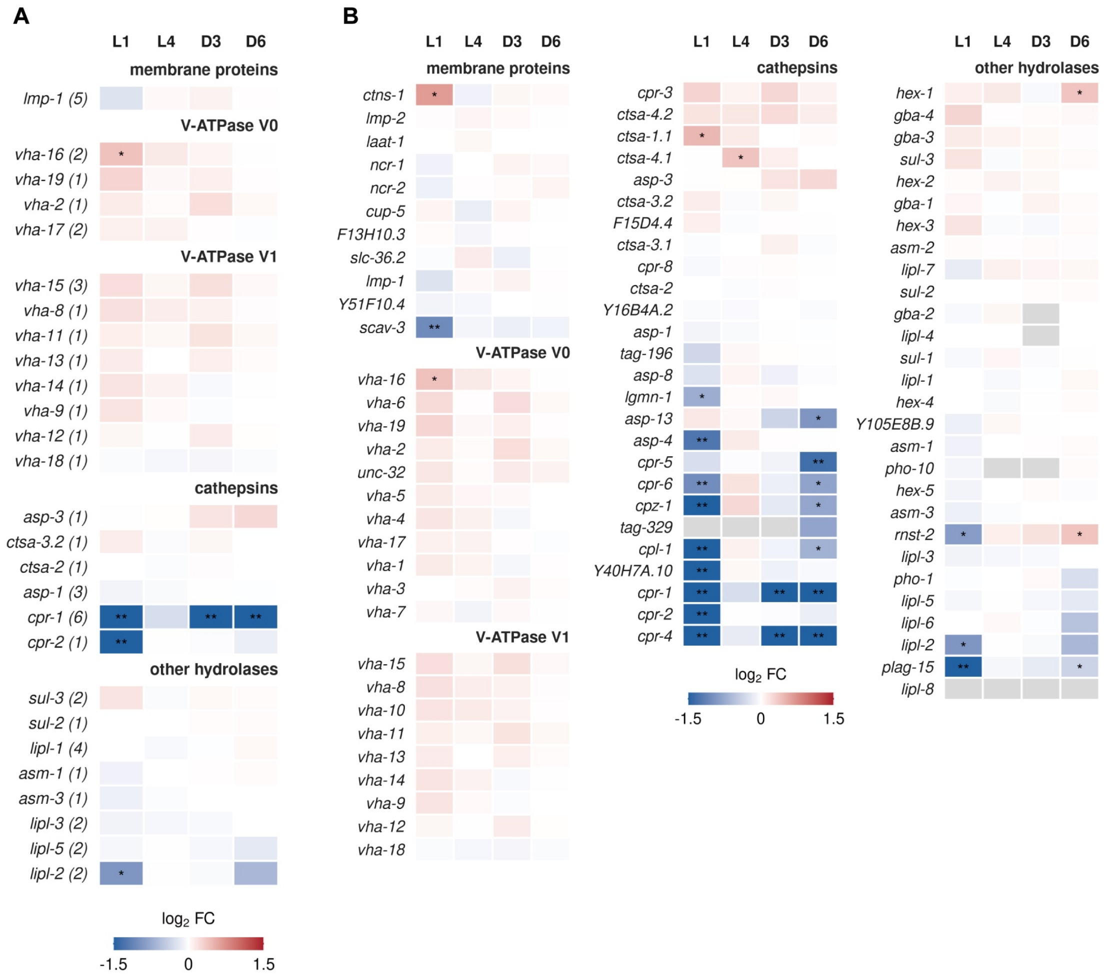
A
L1 L4 D3 D6
membrane proteins
lmp-1 (5)
V-ATPase V0
vha-16 (2)
vha-19 (1)
vha-2 (1)
vha-17 (2)
V-ATPase V1
vha-15 (3)
vha-8 (1)
vha-11 (1)
vha-13 (1)
vha-14 (1)
vha-9 (1)
vha-12 (1)
vha-18 (1)
cathepsins
asp-3 (1)
ctsa-3.2 (1)
ctsa-2 (1)
asp-1 (3)
cpr-1 (6)
cpr-2 (1)
other hydrolases
sul-3 (2)
sul-2 (1)
lipl-1 (4)
asm-1 (1)
asm-3 (1)
lipl-3 (2)
lipl-5 (2)
lipl-2 (2)
log2 FC
-1.5
0
1.5
B
L1 L4 D3 D6
membrane proteins
ctns-1
lmp-2
laat-1
ncr-1
ncr-2
cup-5
F13H10.3
slc-36.2
lmp-1
Y51F10.4
scav-3
V-ATPase V0
vha-16
vha-6
vha-19
vha-2
unc-32
vha-5
vha-4
vha-17
vha-1
vha-3
vha-7
V-ATPase V1
vha-15
vha-8
vha-10
vha-11
vha-13
vha-14
vha-9
vha-12
vha-18
cathepsins
cpr-3
ctsa-4.2
ctsa-1.1
ctsa-4.1
asp-3
ctsa-3.2
F15D4.4
ctsa-3.1
cpr-8
ctsa-2
Y16B4A.2
asp-1
tag-196
asp-8
lgmn-1
asp-13
asp-4
cpr-5
cpr-6
cpz-1
tag-329
cpl-1
Y40H7A.10
cpr-1
cpr-2
cpr-4
log2 FC
-1.5
0
1.5
other hydrolases
hex-1
gba-4
gba-3
sul-3
hex-2
gba-1
hex-3
asm-2
lipl-7
sul-2
gba-2
lipl-4
sul-1
lipl-1
hex-4
Y105E8B.9
asm-1
pho-10
hex-5
asm-3
rnst-2
lipl-3
pho-1
lipl-5
lipl-6
lipl-2
plag-15
lipl-8

**Fig. S9. HLH-30 binds the promoters of genes down-regulated in the *hlh-30* mutant more often than chance at L1 and Day 3.** (A) Percentage of down-regulated (blue) and up-regulated (red) DEGs in *hlh-30(tm1978)* vs WT at each stage (D3, Day 3; D6, Day 6) that have an HLH-30-bound promoter. A promoter was called bound when a peak summit found in both L4 replicates (summits within 100 bp) lay within −1,000 to +500 bp of the TSS. Numbers give bound DEGs over all DEGs. Error bars show Wilson 95% confidence intervals. The dashed line marks the bound share among all other tested genes. *$q < 0.05$, ***$q < 0.001$, two-sided Fisher's exact test against all other tested genes at that stage with Benjamini-Hochberg (BH) adjustment. (B) Pairwise differences in the bound share between stages, for down-regulated (top) and up-regulated (bottom) DEGs. The percentage under each stage is the bound share from (A), and color shows the difference in percentage points (column minus row) for significant pairs. A gene can be a DEG at more than one stage, so each pair was tested with an exact permutation test that allows for the shared genes, BH-adjusted within each direction. *$q < 0.05$, ***$q < 0.001$. The *P* value in each title is the overall test across the four stages (permutation test, 20,000 permutations).

**S9**

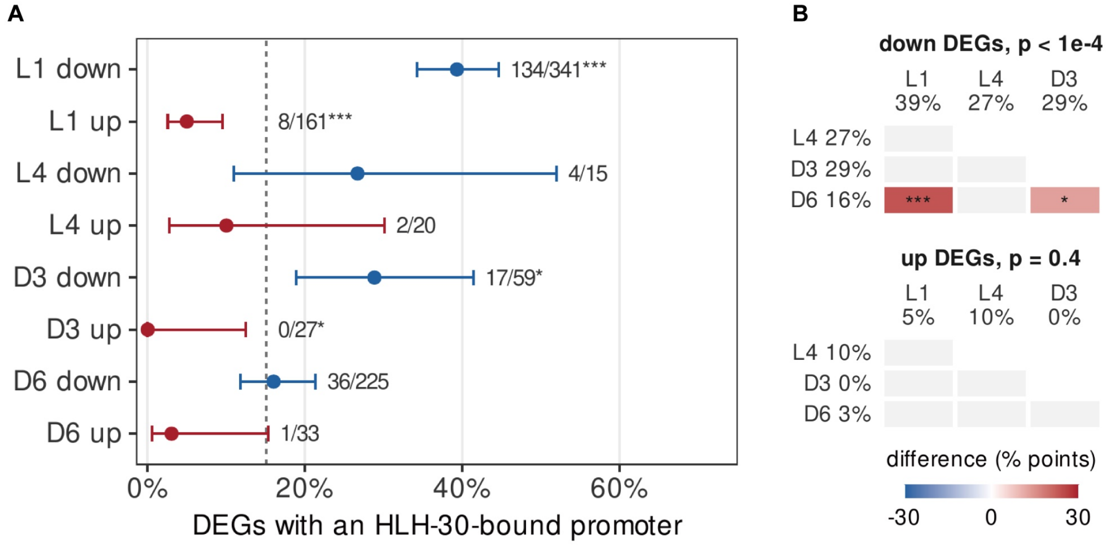
A
L1 down
L1 up
L4 down
L4 up
D3 down
D3 up
D6 down
D6 up
134/341***
8/161***
4/15
2/20
17/59*
0/27*
36/225
1/33
0%
20%
40%
60%
DEGs with an HLH-30-bound promoter
B
down DEGs, p < 1e-4
L1 39%
L4 27%
D3 29%
L4 27%
D3 29%
D6 16%
***
*
up DEGs, p = 0.4
L1 5%
L4 10%
D3 0%
L4 10%
D3 0%
D6 3%
difference (% points)
-30
0
30

**Fig. S10. Promoters of genes differentially expressed in the *hlh-30* mutant are enriched for E-box and other transcription factor motifs.** Simple Enrichment Analysis (SEA, MEME Suite) of all nematode motifs in JASPAR 2022 and all *C. elegans* motifs in CIS-BP 2.00, in the 500 bp (left) and 2,000 bp (right) upstream of the TSS. Columns are down- and up-regulated DEGs in *hlh-30(tm1978)* vs WT at each stage (D3, Day 3; D6, Day 6), and the DEGs of any stage (any down, any up). Controls were the promoters of genes with adjusted $P \geq 0.05$ at that stage, or at every stage for the pooled columns. Rows are the 25 transcription factors with the smallest $q$ among those with a motif enriched ($q < 0.05$) in at least one gene set at a stage where the factor is expressed. All enriched motifs are listed in Supplementary File 6. When a factor has more than one motif, the cell shows the motif with the smallest $q$. E-box, the motif consensus holds an E-box core (CANNTG). inf., CIS-BP motif inferred from a related DNA-binding domain. Color shows fold enrichment over the control. *$q < 0.05$, **$q < 0.01$, ***$q < 0.001$. Hatched cells, the factor is not expressed at that stage in our data.

**S10**

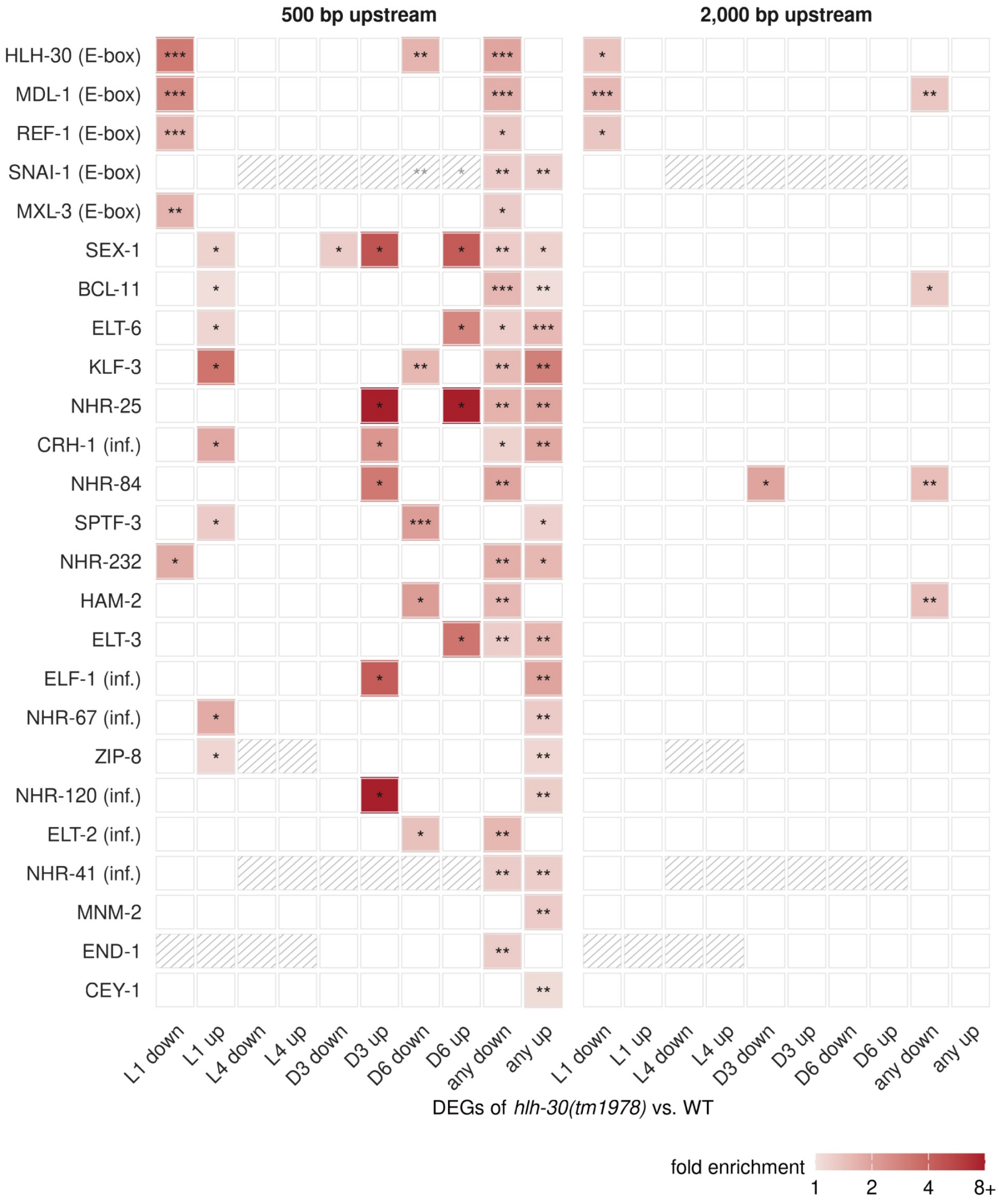
500 bp upstream
2,000 bp upstream
HLH-30 (E-box)
MDL-1 (E-box)
REF-1 (E-box)
SNAI-1 (E-box)
MXL-3 (E-box)
SEX-1
BCL-11
ELT-6
KLF-3
NHR-25
CRH-1 (inf.)
NHR-84
SPTF-3
NHR-232
HAM-2
ELT-3
ELF-1 (inf.)
NHR-67 (inf.)
ZIP-8
NHR-120 (inf.)
ELT-2 (inf.)
NHR-41 (inf.)
MNM-2
END-1
CEY-1
L1 down
L1 up
L4 down
L4 up
D3 down
D3 up
D6 down
D6 up
any down
any up
DEGs of *hlh-30(tm1978)* vs. WT
fold enrichment
1
2
4
8+

**Fig. S11. Genes differentially expressed in the *hlh-30* mutant show largely concordant overlap with published *hlh-30*-dependent gene lists.** Rows are published gene lists in four groups: (1) *hlh-30* loss vs WT in whole animals, (2) *hlh-30* loss in a modified background or one tissue, (3) genes induced by a stimulus in WT that need *hlh-30* and (4) evidence of HLH-30 binding, motif or protein. Numbers next to each list give its size. In groups 1 and 2, triangles give the direction of the published change in the *hlh-30* sample (down, blue; up, red). Group 3 has no triangles because each list holds genes that need *hlh-30* for their induction. The two lists marked NOT *hlh-30*-dependent are genes that *S. aureus* induced in WT without the need for *hlh-30*, shown for comparison. Columns are our down-regulated (blue) and up-regulated (red) DEGs in *hlh-30(tm1978)* vs WT at each stage (D3, Day 3; D6, Day 6). The number below each column label is the percentage of all tested genes that are DEGs in that direction at that stage (the chance rate). Each cell gives the percentage of the published list that is one of our DEGs, and is shown only when the overlap differs from chance (two-sided Fisher's exact test with Benjamini-Hochberg adjustment over all cells, $q < 0.05$). Color shows the fold over chance, red for more overlap than chance and blue for less. The universe at each stage was the genes with an adjusted $P$ value from DESeq2 (tested genes), and genes without one were removed from the published lists.

**S11**

| | n | | L1 ▼ | L1 ▲ | L4 ▼ | L4 ▲ | D3 ▼ | D3 ▲ | D6 ▼ | D6 ▲ |
|---|---|---|---|---|---|---|---|---|---|---|
| | | | 2 | 0.9 | 0.1 | 0.1 | 0.4 | 0.2 | 1 | 0.2 |
| ***hlh-30* loss vs WT, whole animal** | | | | | | | | | | |
| Grove 2009: embryo-L2, fed | 121 | ▼ | 45 | | 3 | | 6 | | 11 | |
| | 11 | ▲ | | | | 9 | | 18 | | |
| Gerisch 2020: fed adults | 108 | ▼ | 36 | | 7 | | 14 | | 28 | |
| | 11 | ▲ | | 20 | | 10 | | 29 | | |
| Gerisch 2020: starved adults | 1,795 | ▼ | 11 | 0.4 | 0.5 | | 2 | | 5 | |
| | 1,660 | ▲ | 0.1 | 3 | | 0.6 | | 1.0 | 2 | 0.6 |
| Munoz-Barrera 2026: starved L1 | 3,841 | ▼ | 7 | <0.1 | | | 1.0 | | 2 | |
| | 3,820 | ▲ | 0.5 | 3 | | 0.3 | <0.1 | | | 0.3 |
| Murphy 2019: starved L1 | 77 | ▼ | | | | | | | | |
| | 52 | ▲ | | | | | | | | |
| Sohn 2025: fed adult males | 304 | ▼ | 26 | | 3 | | 9 | | 17 | 1.0 |
| ***hlh-30* loss in a modified background or one tissue** | | | | | | | | | | |
| Shioda 2023: neuronal *hlh-30* RNAi, glp-1 | 70 | ▼ | 57 | | 7 | | 14 | | 31 | |
| | 23 | ▲ | | | | | | | | |
| Lin 2024: Abeta strain GMC101 | 12 | ▼ | | | | | | | 17 | |
| | 45 | ▲ | 11 | | | | | | | |
| Metcalf 2024: glial XBP-1s | 13 | ▼ | 62 | | 31 | | 38 | | 38 | |
| | 24 | ▲ | | 12 | | | | 8 | 12 | |
| **Induced by a stimulus in WT, needs *hlh-30*** | | | | | | | | | | |
| Visvikis 2014: *S. aureus* | 628 | | 17 | | 0.6 | | 2 | | 6 | |
| Wani 2021: *S. aureus* vs starved | 98 | | | | | 2 | | | | 2 |
| Chen 2017: Cry5B toxin | 516 | | 8 | | | | | | 4 | |
| Hsieh 2026: Cry5B toxin (HLH-30 R38K) | 280 | | | | | | | | | |
| Possik 2023: pgph-2 overexpression | 206 | | 7 | | 2 | | 5 | | 9 | |
| Murphy 2019: starved L1 | 53 | | 13 | | | | | | | |
| Visvikis 2014: *S. aureus*, NOT *hlh-30*-dependent | 184 | | 5 | 4 | 2 | | 4 | | 9 | |
| Wani 2021: *S. aureus*, NOT *hlh-30*-dependent | 6 | | | | | | | | | |
| **Binding, motif or protein evidence** | | | | | | | | | | |
| ENCODE: ChIP, L4 | 3,910 | | 4 | 0.5 | | | | <0.1 | | |
| ENCODE: ChIP, embryo | 1,303 | | 6 | 0.2 | | | <0.1 | | 0.5 | |
| Grove 2009: HLH-30 binding (PBM) | 404 | | 6 | | | | | | | |
| Harvald 2017: CLEAR motif, starvation | 487 | | 6 | | | | 1 | | 4 | |

vs chance: 16x fewer — 4x fewer — chance — 4x more — 16x more
