## Additional file 9 - Supp methods for "*Caenorhabditis elegans* HLH-30/TFEB has age-dependent effects on the transcriptome and is required for normal progeny number and larval survival under basal growth conditions"

### Supplementary Methods: RNA-seq data processing and bioinformatic analyses

#### RNA sequencing

Libraries were sequenced by Plasmidsaurus (3'-end RNA-seq, single-end reads with unique molecular identifiers).

#### Read processing, mapping and quantification

Read processing up to deduplicated alignments was performed by Plasmidsaurus. Read quality was assessed with FastQC v0.12.1 (Andrews, 2010). Reads were filtered with fastp v0.24.0 (Chen et al., 2018; Chen, 2023) with poly-X tail trimming, 3' quality-based tail trimming, a minimum Phred quality of 15 and a minimum length of 25 bp. Filtered reads were aligned to the *C. elegans* reference genome (WBcel235; Plasmidsaurus build WBcel235_62) with STAR v2.7.11b (Dobin et al., 2013) with removal of non-canonical splice junctions, and alignments were coordinate-sorted with samtools (Danecek et al., 2021; v1.21 for the L1 libraries and v1.23 for the L4 to day 6 libraries, as recorded in the BAM headers). PCR and optical duplicates were removed by UMI-based deduplication with UMIcollapse v1.1.0 (Liu, 2019). Alignment quality, strand specificity and read distribution were assessed with RSeQC v5.0.4 (Wang et al., 2012) and Qualimap v2.3 (Okonechnikov et al., 2016), and summarized with MultiQC v1.33 (Ewels et al., 2016).

Gene-level counts were generated from the deduplicated alignments with featureCounts (Liao et al., 2014) in Rsubread v2.22.1 (Liao et al., 2019), using exon features of the Ensembl release 115 annotation of WBcel235 (Dyer et al., 2025) summarized by gene. Reads were counted forward-stranded, as 3'-end library reads match the sense strand of the transcript. Reads that mapped to several loci were counted at each locus (Rsubread default). Genes with fewer than 10 reads summed over all samples of an analysis were removed.

#### Removal of unwanted variation and differential expression analysis

Two sample sets of wild type and *hlh-30(tm1978)* were analyzed, the L4 to day 6 set (28 samples) and the full set (34 samples), which also included L1 larvae. Each set had three biological replicates per genotype at L1 and L4, six at day 3 of adulthood and five at day 6 of adulthood. The adult replicates came from two cohorts, one grown at room temperature (22 °C) and one at 20 °C (three replicates per genotype from each cohort at day 3, and two and three at day 6). When growth temperature was added to the model, 91% to 98% of the DEGs at each stage were still DEGs, and the two cohorts intermixed in the PCA (Fig. S1C). Hence, the two cohorts were pooled as one group to increase statistical power. A day 3 and day 6 subset (22 samples) was used for the adult-only PCA and clustering.

For each set, a single factor of unwanted variation (W_1) was estimated with RUVSeq v1.42.0 (Risso et al., 2014), using the RUVs method (k = 1) with the replicate samples of each genotype × stage group as negative-control sample sets. RUVs was applied to all filtered genes after upper-quartile normalization (EDASeq `betweenLaneNormalization`; Bullard et al., 2010).

Differential expression was tested with DESeq2 v1.48.2 (Love et al., 2014) in R v4.5.2 (R Core Team, 2025) with the design `~ W_1 + replicate + condition`, where `condition` combines genotype and stage and `replicate` is the biological-replicate number. Wald tests were run for every pair of conditions that share a genotype (developmental transitions) or a stage (mutant vs wild type). Log2 fold changes were shrunk with ashr v2.2.63 (Stephens, 2017). P values were adjusted with the Benjamini-Hochberg method (Benjamini and Hochberg, 1995) after DESeq2's default independent filtering. Genes with an adjusted $P < 0.05$ and an absolute shrunken log2 fold change $\geq 1$ were called differentially expressed genes (DEGs).

For visualization and clustering, counts were variance-stabilized with DESeq2 (`vst`, blind = FALSE). The replicate effect and W_1 were then removed with limma v3.64.3 `removeBatchEffect` (Ritchie et al., 2015), with the condition design supplied so that genotype and stage differences were kept ("corrected VST"). The corrected VST was not used for differential expression.

#### Principal component analysis

Principal component analysis was performed with R `prcomp` on the corrected VST values of the 500 most variable genes, with each gene centered and scaled to unit variance. For the L1-only analysis (six samples), counts were variance-stabilized without a design (`vst`, blind = TRUE) and no correction was applied.

#### Hierarchical clustering and heat map

For the sample correlation heat maps, corrected VST values were first averaged over the replicates of each genotype × stage group. The 1,000 genes with the highest variance across the group means were selected, and Pearson correlation coefficients were computed between groups. Groups were ordered by hierarchical clustering with average linkage (Sokal and Michener, 1958) on the Euclidean distances between rows of the correlation matrix. Heat maps were drawn with ComplexHeatmap v2.26.1 (Gu et al., 2016).

#### Gene clustering (k-means and HDBSCAN)

The genes clustered in each sample set were the DEGs of any contrast of that set. Corrected VST values were averaged per genotype × stage condition and z-scored per gene across conditions.

k-means clustering used R `kmeans` (Hartigan-Wong algorithm; Hartigan and Wong, 1979) with 25 random starts, at most 100 iterations and a fixed random seed. The number of clusters was chosen by the elbow method (Thorndike, 1953). k-means was run for $k = 2$ to 12, and the chosen k was the point of the within-cluster sum of squares curve (scaled from 0 to 1) that was farthest from the straight line between the two ends of the curve. This gave k = 6 for both the L4 to day 6 set and the full set. For the day 3 and day 6 subset, k was set to 4.

As a density-based comparison, genes were also clustered with HDBSCAN (Campello et al., 2013) in the R package dbscan v1.2.4 (Hahsler et al., 2019), with a minimum cluster size of 5 genes and Euclidean distance. For HDBSCAN, each gene was z-scored across samples and then averaged per condition. Genes that HDBSCAN did not assign to a cluster were reported as noise.

Gene clusters were displayed with UMAP (McInnes et al., 2018) in the R package uwot v0.2.4 (Melville, 2025) with 15 nearest neighbors, a minimum distance of 0.1, Euclidean distance and a fixed random seed.

**Gene Ontology analysis**

Gene Ontology (GO) enrichment (Ashburner et al., 2000; Gene Ontology Consortium, 2023) was tested with clusterProfiler v4.16.0 `enrichGO` (Yu et al., 2012; Wu et al., 2021) and the org.Ce.eg.db v3.21.0 annotation (Carlson, 2025) for biological process, cellular component and molecular function. WormBase gene identifiers (Sternberg et al., 2024) were mapped to Entrez identifiers, and unmapped genes were dropped. The background was all genes that passed the count filter in the same analysis. Gene sets of 10 to 500 genes were tested, and terms with a Benjamini-Hochberg adjusted $P < 0.05$ (and $q < 0.2$) were considered enriched. Gene lists tested were the k-means clusters, the up- and down-regulated DEGs of each *hlh-30(tm1978)* vs wild-type contrast and the gene lists from the comparison of developmental changes between genotypes (next section). The terms shown in the figures were selected from the enriched terms of each list.

**Comparison of developmental expression changes between genotypes**

To find genotype-specific developmental changes, the DEGs of each developmental transition (L1 to L4, L4 to day 3, L4 to day 6 and day 3 to day 6) were compared between wild type and *hlh-30(tm1978)*. Only genes that DESeq2 tested in both genotypes' contrasts were compared, so that a gene removed by independent filtering in one genotype was not called exclusive to the other. A gene was "exclusive" when it was a DEG in only one genotype, "shared" when it was a DEG in both genotypes with an absolute difference in log2 fold change < 1, and "differential" when it was a DEG in both genotypes with an absolute difference ≥ 1. Exclusive genes were split by the direction of change. Differential genes that changed in the same direction in both genotypes were called "more up" or "more down" in the genotype with the larger change, and those that changed in opposite directions were called "opposite".

Two gene sets were built for Fig. S3 from the GO terms enriched (adjusted $P < 0.05$) in any gene category (above) of the three adult transitions. The oocyte and germline set (Fig. S3A and S3C) was the union of the genes in every such term whose name contains a germline-related word (germ, gamete, oogenesis, oocyte, meiosis, P granule, reproduction, gonad, fertilization or sex determination) and no sperm-related word (sperm, male gamete or pseudopodium), 31 genes. The sperm set (Fig. S3B and S3D) was the 18 genes of the enriched pseudopodium term, which in *C. elegans* holds the major sperm protein genes. For display, the mean variance-stabilized expression of each gene in each genotype and stage of the four-stage analysis was z-scored across the eight group means. For each set, the difference between genotypes at each stage was tested with limma camera (Wu and Smyth, 2012) on the corrected VST, which accounts for the correlation between the genes of the set. Camera P values were adjusted with the Benjamini-Hochberg method over the eight tests (two sets, four stages). Single genes were tested with the DESeq2 mutant vs wild-type contrast of that stage.

**Analysis of modENCODE's HLH-30 ChIP-seq data**

HLH-30::GFP ChIP-seq data from the modENCODE project (Snyder laboratory; Kudron et al., 2018) were used for L4 larvae (GEO GSE48751; SRX331310 and SRX331312, inputs SRX331309 and SRX331311) and late embryos (GEO GSE48752; SRX331314 and SRX331316, inputs SRX331313 and SRX331315). Peaks (MACS2, Q < 1e-5; Zhang et al., 2008) and coverage tracks (bigWig) aligned to ce11 were obtained from ChIP-Atlas (Oki et al., 2018; Zou et al., 2024). Gene coordinates were taken from the Ensembl release 115 annotation, and chromosome names were converted to UCSC ce11 names.

For the metagene profile, coverage was summed in 60 bins over ±3 kb around the transcription start sites (TSS) of protein-coding genes (a random sample of up to 20,000 TSS), with minus-strand genes reversed. Each track was divided by its genome-wide mean, ChIP signal was divided by the matching input signal and the profile was scaled so that the flanks beyond ±2.5 kb equal 1. Locus tracks used genome-mean-normalized coverage in 800 bins over ±3 kb around the TSS. An L4 peak was taken as reproducible when the summit of a peak in replicate SRX331310 was within 100 bp of a summit in replicate SRX331312.

**Motif analysis**

Promoters were defined as 2,000 bp upstream to 200 bp downstream of each gene's TSS and extracted from BSgenome.Celegans.UCSC.ce11 v1.4.2. Sites of the HLH-30 position weight matrix from JASPAR (MA1449.1; Rauluseviciute et al., 2024) were found with motifmatchr v1.32.0 (Schep, 2025), which uses MOODS (Korhonen et al., 2009), and TFBSTools v1.48.0 (Tan and Lenhard, 2016), at a match P < 1e-4 with the base composition of the scanned promoters as background. Sites therefore include close variants of the consensus TCACGTGA (such as TCATGTGA). A gene with at least one site counted as a motif-containing gene. For each of the 12 DEG lists (all, up and down DEGs of the mutant vs wild-type contrast at L1, L4, day 3 and day 6), the fraction of motif-containing genes was compared with that of the other expressed genes (the 18,597 genes that passed DESeq2's independent filtering in at least one mutant vs wild-type contrast) by a two-sided Fisher's exact test, with Benjamini-Hochberg correction over the 12 tests. In the promoter schematics, E-box sites are exact CACGTG matches on either strand (the CLEAR element core; Sardiello et al., 2009). For the *tts-1*, *cpr-4* and *ztf-28* promoters, three more elements were counted in addition to the JASPAR HLH-30 sites: the E-box (CACGTG), the DAF-16-binding element (DBE; TRTTTAC and TTGTTTAC) and the DAF-16-associated element (DAE; TGTAAG and TGATAAG), which is bound by PQM-1. The DBE and DAE were included because they are enriched at HLH-30 binding sites (Lin et al., 2018).

**Motif analyses to identify transcription factors that may contribute to the differential gene expression in the *hlh-30(tm1978)* mutant**

Motif enrichment in DEG promoters was tested as in Chang et al. (2026) with SEA (Bailey and Grant, 2021) in the MEME Suite v5.5.7 (Bailey et al., 2015), with all motifs of JASPAR 2022 CORE nematodes (non-redundant; Castro-Mondragon et al., 2022) and CIS-BP 2.00 *C. elegans* (Weirauch et al., 2014). Sequences were the 500 bp and the 2,000 bp upstream of each TSS, tested as two separate analyses. The DEGs of the mutant vs wild-type contrast in the full sample set were tested for each stage and direction. The control sequences were the promoters of the genes tested at that stage with adjusted $P \geq 0.05$. The DEGs of all stages were also pooled for each direction, with the genes that were tested at every stage and were never a DEG as control sequences. Gene sets with fewer than 10 genes were not tested. Motifs with SEA $q < 0.05$ were considered enriched. A motif was kept only when the gene of its transcription factor was expressed at that stage (mean DESeq2-normalized count ≥ 10 over the samples of that stage, or of any stage for the pooled sets). The figure shows the 25 transcription factors with the smallest q, each with its best motif. CIS-BP motifs inferred from a related DNA-binding domain were marked, and motifs with an E-box core (CANNTG) in their consensus were grouped.

**HLH-30 binding and published HLH-30 gene lists**

These analyses (Fig. S7 to S9 and S11) used the DEGs of the mutant vs wild-type contrast at each stage in the full sample set. The genes compared at a stage were those that passed DESeq2's independent filtering at that stage.

A gene was counted as bound by HLH-30 when a reproducible L4 summit (section "Analysis of HLH-30 ChIP-seq data") was within 1,000 bp upstream to 500 bp downstream of its TSS. For each stage and direction, the bound share of the DEGs was compared with the bound share of the other genes by a two-sided Fisher's exact test, with Benjamini-Hochberg correction over the eight tests, and shown with a Wilson 95% confidence interval (Wilson, 1927). The bound shares of two stages' DEG lists of the same direction were compared by an exact permutation test instead of a Fisher's exact test, because a gene that is a DEG at both stages is in both lists. The genes were split into three groups (first list only, both lists and second list only). The P value was the probability that a random spread of the bound genes over these groups gives a difference in bound share at least as large as observed, in either direction. P values were adjusted with the Benjamini-Hochberg method over the six stage pairs of each direction. The overall difference across the four stages was tested for each direction with the chi-squared statistic of the stage by binding table, in which a gene counts at every stage where it is a DEG. Its P value came from 20,000 permutations of the binding labels among the genes of that direction.

Published HLH-30 gene lists were taken from the supplementary tables of each study and mapped to WormBase gene identifiers by name. When a study gave only a full table, we applied a cutoff, which is listed with the gene lists in the code repository. The lists were in four groups, (1) genes lower or higher after loss of *hlh-30* in whole animals (Grove et al., 2009; Murphy et al., 2019; Gerisch et al., 2020; Sohn et al., 2025; Muñoz-Barrera et al., 2026), (2) the same comparison in a modified background or after loss in one tissue (Shioda et al., 2023; Lin et al., 2024; Metcalf et al., 2024), (3) genes induced by a stimulus in wild type that need *hlh-30* for the induction (Visvikis et al., 2014; Chen et al., 2017; Murphy et al., 2019; Wani et al., 2021; Possik et al., 2023; Hsieh et al., 2026), with the *hlh-30*-independent induced genes of two of these studies as controls (Visvikis et al., 2014; Wani et al., 2021), and (4) binding, motif or protein evidence, namely the L4 and late-embryo ChIP-seq data above (summit within 1,000 bp upstream to 200 bp downstream of a TSS), targets predicted from protein-binding microarray data (Grove et al., 2009) and starvation-induced proteins with a CLEAR motif (Harvald et al., 2017). For each list, stage and direction, the overlap with our DEGs was tested by a two-sided Fisher's exact test, using only the genes compared at that stage, with Benjamini-Hochberg correction over all 232 tests. For each overlap with adjusted $P < 0.05$, the figure shows the percentage of the list that is one of our DEGs, colored by the fold change over chance (observed overlap divided by expected overlap).

Core autophagy genes were the 69 genes of Table 1 in Zhang and Meléndez (2025), and core lysosome genes were the 85 genes of Supplementary file 1 in Sun et al. (2020), in their five categories. Gene names were mapped as above, except *ctsa-1*, which was mapped to C08H9.1 (now *ctsa-3.2*), the gene it named in Sun et al. (2020), because WormBase now uses *ctsa-1* as an alias of K10B2.2, a different gene that the table also lists. Genes linked to *hlh-30* were those reported in at least one published study to have lower mRNA or reporter expression, or a lost or reduced induction, in an *hlh-30* mutant or after *hlh-30* RNAi (studies listed in Supplementary File 5). For each gene and stage, the figures show the log2 fold change of the mutant vs wild-type contrast, with marks for adjusted $P < 0.05$ and for a DEG.

**Software**

| Step | Software (version) |
|---|---|
| Read processing and alignment (Plasmidsaurus) | FastQC 0.12.1, fastp 0.24.0, STAR 2.7.11b, samtools 1.21 / 1.23, UMIcollapse 1.1.0, RSeQC 5.0.4, Qualimap 2.3, MultiQC 1.33 |
| Pipeline (R 4.5.2, Bioconductor 3.21; Docker image rayandrew/celegans-r-analysis:0.11.0) | Rsubread 2.22.1, RUVSeq 1.42.0, EDASeq 2.42.0, DESeq2 1.48.2, ashr 2.2.63, limma 3.64.3, clusterProfiler 4.16.0, org.Ce.eg.db 3.21.0, dbscan 1.2.4, uwot 0.2.4, cluster 2.1.8.1 |
| Figures and post-hoc analyses (R 4.5.2, Bioconductor 3.22) | ComplexHeatmap 2.26.1, ggplot2 4.0.2, uwot 0.2.4, motifmatchr 1.32.0, TFBSTools |

| Step | Software (version) |
| --- | --- |
| | 1.48.0, JASPAR2024 0.99.7, BSgenome.Celegans.UCSC.ce11 1.4.2, GenomicRanges 1.62.1, rtracklayer 1.70.1 |
| Motif analyses | MEME Suite 5.5.7 (SEA) |

**Data and code availability**

The RNA-seq data generated in this study (raw reads and gene counts) are available in the NCBI Gene Expression Omnibus (GEO) under accession GSE354145. Analysis code is available at https://github.com/sherlynpw/hlh30-rnaseq and is archived on Zenodo at https://doi.org/10.5281/zenodo.24213667.
